# Evolution-driven attenuation of alphaviruses highlights key glycoprotein determinants regulating viral infectivity and dissemination

**DOI:** 10.1101/636779

**Authors:** Maria G. Noval, Bruno A. Rodriguez-Rodriguez, Margarita Rangel, Kenneth A. Stapleford

## Abstract

Understanding the fundamental mechanisms of arbovirus transmission and pathogenesis is essential to develop new strategies for treatment and prevention. We previously took an *in vivo* evolution-based approach and identified the chikungunya virus E1 glycoprotein residue 80 to play a critical role in viral transmission and pathogenesis. In this study, we address the genetic robustness and function of position 80 and demonstrate that this highly conserved residue is a key determinant in alphavirus infectivity and dissemination through modulation of viral fusion and cholesterol dependence. In addition, in studying the evolution of position 80, we identified a network of glycoprotein residues, including epidemic determinants, that regulate virus dissemination and infectivity. These studies underscore the importance of taking evolution-based approaches to not only identify key viral determinants driving arbovirus transmission and pathogenesis but also to uncover fundamental aspects of arbovirus biology.

## INTRODUCTION

Arboviruses (arthropod-borne viruses) represent a significant global health threat capable of causing severe disease in humans and substantial economic burden worldwide (Labeaud et al., 2011; Rodriguez-Morales et al., 2016). During the past several decades, arboviruses such as chikungunya virus (CHIKV), West Nile Virus (WNV), Zika virus (ZIKV), yellow fever virus, and dengue virus have caused devastating outbreaks, spreading to naïve populations and endogenous vectors (Gould et al., 2017; Vignuzzi and Higgs, 2017). This global expansion, and the absence of effective antivirals and vaccines, (Powers, 2018; Silva and Dermody, 2017; Wilder-Smith et al., 2017) points towards an urgent need to study the fundamental aspects of arbovirus biology which are likely to provide avenues for the development of new antiviral therapies.

Viral evolution is a key driving force for arbovirus emergence and in turn can be used to study how arboviruses are transmitted and cause disease. Recent natural outbreaks of Venezuelan equine encephalitis virus, WNV, ZIKV, and CHIKV have been retrospectively mapped to adaptive changes in the viral glycoproteins, which have led to increases in transmission, pathogenesis (Ebel et al., 2004; Greene et al., 2005; Moudy et al., 2007; Pesko and Ebel, 2012; Yuan et al., 2017), and the adaptation to new vectors (Brault et al., 2004; Tsetsarkin et al., 2007). Although the viral glycoproteins are not the only factors affecting viral transmission, extensive studies of natural and laboratory derived glycoprotein variants have demonstrated that the glycoproteins are critical determinants impacting emergence, transmission, pathogenesis, and arboviral disease (Brault et al., 2004; Greene et al., 2005; Stapleford et al., 2014). However, the underlying molecular mechanisms of how these proteins function in arbovirus transmission and infectivity are not completely understood.

To study arbovirus transmission, we use CHIKV, a member of the *Togaviridae* family (genus alphavirus) transmitted by the *Aedes* species of mosquitoes (Higgs and Vanlandingham, 2015). CHIKV is a single-stranded positive-sense RNA virus with a genome that codes for ten protein products, four nonstructural proteins (nsP1-4) and six structural proteins (CP, E3, E2, 6K, TF, and E1). The E1 fusion glycoprotein is a class-II fusion protein (Kielian and Rey, 2006), anchored to the membrane at the C-terminus, and with an ectodomain sub-divided into domains I-III, with domain II containing the fusion loop (Voss et al., 2010). In the mature virion, E1 and the attachment protein E2 are arranged to form 80 trimeric spikes constituted of trimers of E1-E2 heterodimers (Kielian and Rey, 2006; Sun et al., 2013). These protein complexes then mediate CHIKV internalization by receptor mediated endocytosis and fusion within the early endosome (Hoornweg et al., 2016), where low endosomal pH triggers E1-E2 dissociation, E1 fusion loop insertion into the target membrane, and concomitant membrane fusion (Kielian and Rey, 2006; van Duijl-Richter et al., 2015).

The CHIKV glycoproteins play a significant role in CHIKV transmission, emergence, and spread. In particular, an adaptive mutation in the CHIKV E1 glycoprotein, A226V, gave rise to the Indian Ocean Linage (IOL) of CHIKV and represents one of the most emblematic examples of the role of viral glycoproteins in transmission and adaptation (Schuffenecker et al., 2006). The emergence of the E1-A226V mutation was found to increase transmission by *Ae. albopictus* mosquitoes conferring a selective advantage over the vector *Ae. aegypti* (Tsetsarkin et al., 2007; Vazeille et al., 2007). As a consequence, this mutation allowed for rapid spread from the Indian Ocean, to India, Italy and eventually France, due to its ability to infect and to be transmitted by the widespread *Ae. albopictus* mosquito (Vignuzzi and Higgs, 2017). Since then, CHIKV has continued to evolve, accumulating mutations in the viral attachment glycoprotein E2 (e.g., L210Q, K252Q), which further increased CHIKV fitness in *Ae. albopictus* (Tsetsarkin et al., 2014; Tsetsarkin and Weaver, 2011). This continuous step-wise evolution of CHIKV again highlights the viral glycoproteins as key determinants of arbovirus transmission and infectivity.

In a previous study, we took an *in vivo* evolution-based approach to study novel and emerging viral determinants of CHIKV transmission and pathogenesis (Stapleford et al., 2014). Using this model, we identified a highly transmittable and virulent CHIKV variant in the saliva of both *Ae. aegypti* and *Ae. albopictus* mosquitoes. This variant contained two point mutations in the E1 glycoprotein (E1-V80I and E1-A129V), and we showed that both the double mutant as well as each individual substitution played a critical role in viral transmission and pathogenesis. In particular, position 80 is fully conserved within the alphavirus genus and is located in the structurally conserved tip of domain II (Rey and Lok, 2018; Voss et al., 2010) next to the fusion loop and in topological proximity to the residue E1-226. Given the selection of this variant *in vivo* and its evolutionary conservation, we hypothesize that this residue is a key determinant for CHIKV transmission and infectivity. However, its molecular mechanisms and function remain unknown.

In this study, we aimed to dissect and identify the specific role of the E1 glycoprotein position 80 in alphavirus biology. To be able to fully address the role of this position, we combined a mutational tolerance approach, where we mutated the valine at position 80 to each possible canonical amino acid, with an *in vitro* evolution approach in mammal and mosquito cell lines. Using these approaches, we found that E1 position 80 is a key determinant of alphavirus infectivity and dissemination through the modulation of viral fusion and cholesterol dependence. In addition, we identified a network of genetically linked residues in the E1 glycoprotein, which include novel and previously identified residues involved in viral fusion and cholesterol dependence. Finally, we demonstrate a functional link between E1 position 80 and 226 in mediating viral cholesterol dependence *in vitro* and viral infectivity and dissemination in mammals. Taken together, we show that the alphavirus E1 residue 80 is a key determinant for alphavirus infectivity and dissemination and that this residue functions in concert with other viral residues to orchestrate these processes. Moreover, we demonstrate that coupling mutational tolerance, evolution, and functional analyses delivers new insights into arbovirus transmission mechanisms providing a framework to identify new targets for therapeutic design.

## RESULTS

### Residue 80 in the chikungunya virus E1 glycoprotein tolerates aliphatic amino acids and is genetically constrained in a host-specific manner

We previously identified two variants in the CHIKV E1 glycoprotein, E1-V80I and E1-A129V (Figure 1A), that together increased viral transmission and pathogenesis *in vivo* (Stapleford et al., 2014). Given the selection of E1-V80I *in vivo* and its evolutionary conservation, we hypothesize that this residue is a key determinant for CHIKV transmission and infectivity. To investigate the role of residue E1-80 in the CHIKV life cycle, we began by addressing the mutational tolerance of this position. To do this, we took an unbiased mutagenesis approach and changed position 80 to every possible canonical amino acid residue. Because CHIKV alternates between a mammal host and mosquito vector, we generated each variant in baby hamster kidney (BHK-21) and *Ae. albopictus* (C6/36) cells, passaged the viruses three times over each cell type, and addressed the genetic stability by Sanger sequencing (Table 1).

**Table 1.**
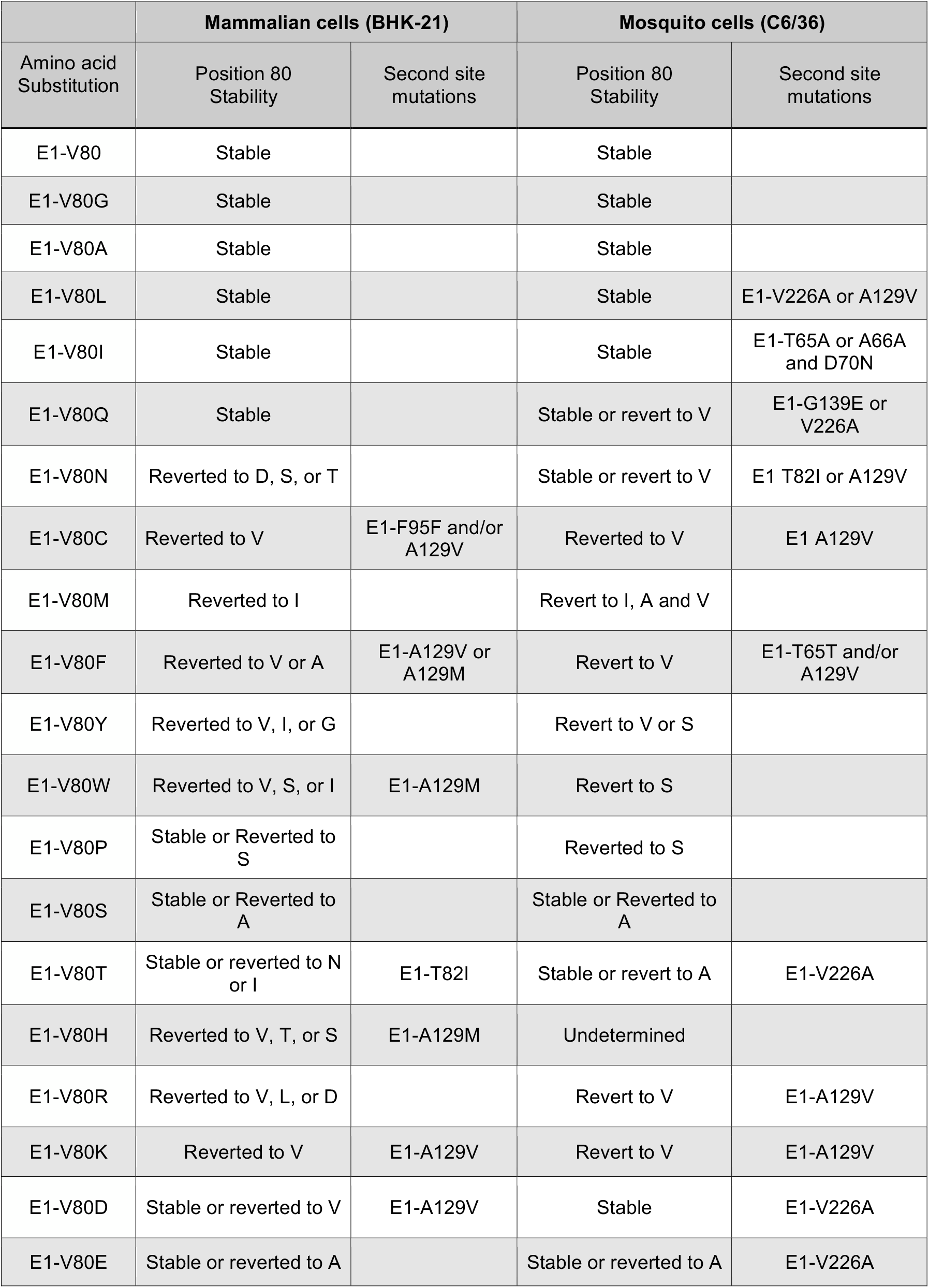
Sequence stability of the E1 V80X variants in mammals and mosquitoes cell lines

**Figure 1.**
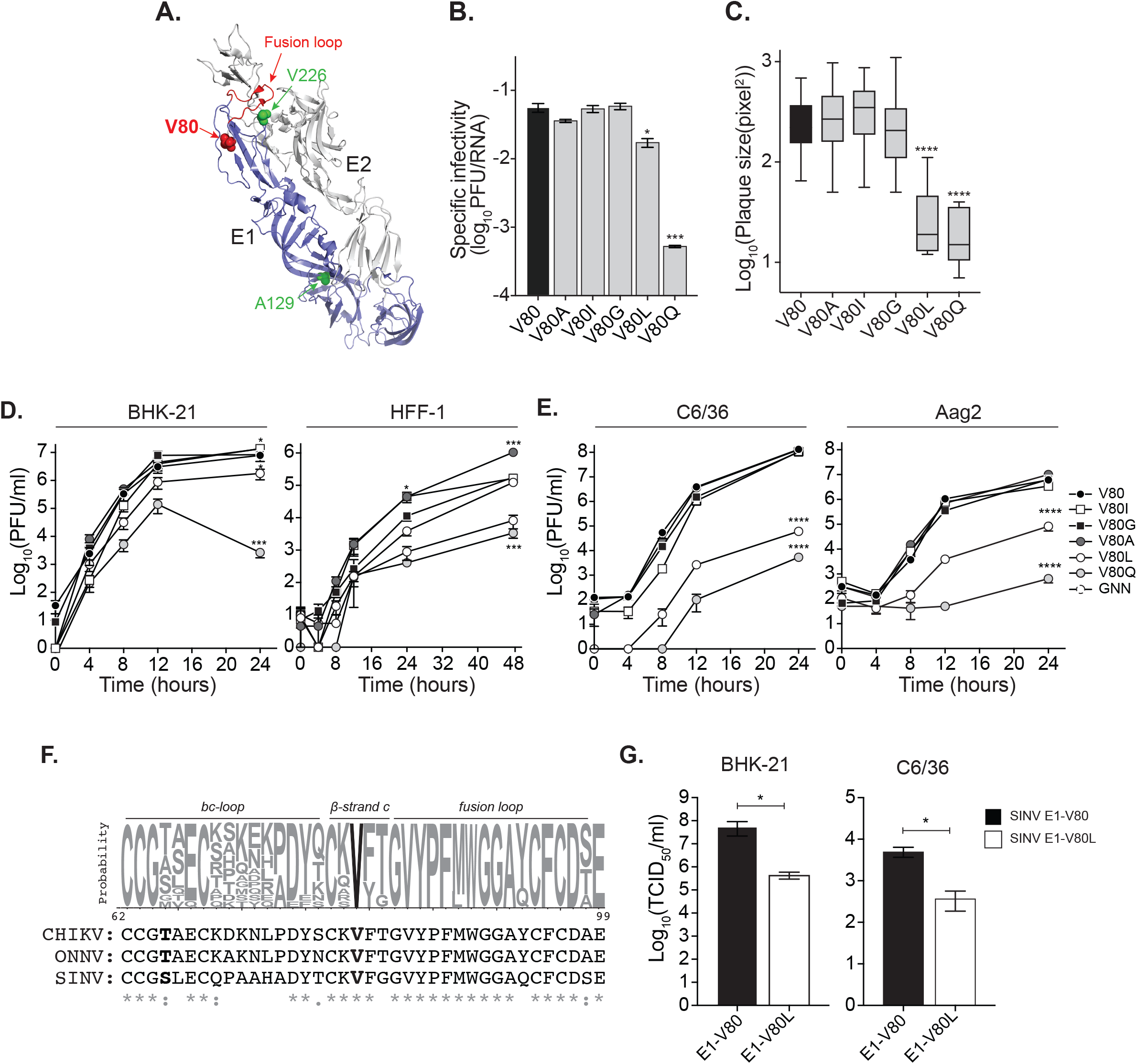
Position 80 of the E1 glycoprotein is a main determinant of alphavirus infectivity *in vitro.* (A) Ribbon representation of the CHIKV E1/E2 heterodimer (PDBID: 3N42) highlighting the E1 monomer (blue) and the E2 monomer (gray) is shown. Residue at position 80 is shown as a red sphere. Positions E1-226 and E1-129 are highlighted in green. The E1-fusion loop, spanning residues 83 to 98, is indicated in red. (B) Specific infectivity expressed as infectious particles over the number of genomes per ml (PFU/RNA) from the BHK-21 stock viruses was determined by plaque assay on Vero cells and by quantification of the number of genomes by RT-qPCR. The bars represent the mean and the standard error of the mean (SEM), N=3 (* p < 0.05, *** p < 0.001; Kruskal-Wallis test). (C) Plaque size in Vero cells was quantified using ImageJ (**** p < 0.0001, N > 40; Kruskal-Wallis test). (D) One-step viral growth kinetics performed in mammal BHK-21 and HFF-1 (E) and mosquito C6/36 and Aag2 (B) cell lines. Cells were infected at a MOI of 0.01 with each variant, and samples of the supernatant were collected at the indicate time points. Results are represented as mean and SEM, N=6 (* p<0.05, *** p<0.001 and **** p<0.0001). P values were determined by two-way ANOVA with Bonferroni post-hoc test. (F) Sequence conservation of position 80 in E1 glycoprotein within the alphavirus genus. Upper panel: Sequence logo showing the probability of each possible amino acid within the region comprising residues 62 to 99 generated from a sequence alignment of 13 representative members of the alphavirus genus (see materials and methods). Positions comprising the *bc-loop, β-strand c* and the fusion loop are indicated. Residue E1-80 is highlighted in bold. Lower panel: CHIKV IOL, ONNV and SINV sequenced alignment. Position E1-80 and E1-65 are highlighted in bold (* = fully conserved residues, : and . = partially conserved residues). (G) BHK-21 cells and C6/36 cells were infected at a MOI of 0.01 with wildtype SINV (E1-V80) or the E1-V80L mutant. Production of infectious particles after 24 hr post infection were determined by TCID_50_. Results are represented as mean and SEM, N=4. P values were determined by Mann-Whitney U test.

In mammalian cells, we found five variants bearing isoleucine (I), glycine (G), alanine (A), leucine (L) and glutamine (Q) substitutions (V80I, G, A, L, and Q) to be genetically stable with the majority of these mutations being aliphatic, similar to the wildtype valine residue (V80). In addition, the unstable mutants did not all revert to wildtype (Table 1 **and Supplemental Figure 1**) but in some cases changed to a stable variant or acquired second-site mutations including E1-A129V which we have previously identified *in vivo* (Stapleford et al., 2014) providing further evidence that these residues are functionally and genetically linked.

On the other hand, in mosquito cells, we found E1-V80G and V80A, to be genetically stable (Table 1 **and Supplementary Figure 1**), as other variants reverted to wildtype or accumulated second-site mutations throughout the E1 glycoprotein. Interestingly, E1-V80L, E1-V80Q, E1-V80D, E1-V80E and E1-V80T remained genetically stable yet acquired the mutation E1-V226A, suggesting a genetic link between these two positions. In addition, we found with the E1-V80I variant, two replicates show second-site mutations: a double mutation E1-A66A:E1-D70N, or a T65A substitution (Table 1 **and Supplementary Figure 1B**). It is interesting to note that position 66, 70 and 226 of E1 glycoprotein have been previously implicated with CHIKV fusion and cholesterol dependence suggesting a potential functional link between position 80 and these residues in these processes (Lu et al., 1999; Tsetsarkin et al., 2011; Vashishtha et al., 1998). Finally, it should be noted that with exception of mutation E1-A129V, none of these mutations were observed in passages performed in mammalian cells (Table 1 **and Supplementary Figure 1)**, suggesting that these second site mutations are host-specific. Taken together, these data suggest host-specific constraints on position E1-80 and that changes at this position are more tolerated in the mammalian host where variants remained genetically stable without the accumulation of second-site mutations. Given this, we chose to focus on the five genetically stable E1-V80 variants generated in mammalian cells (E1-V80I, A, G, L, and Q) for further *in vitro* and *in vivo* studies.

### Chikungunya virus E1-V80 impacts virion production and infectivity *in vitro*

Upon generation of infectious virions in BHK-21 cells, we observed that two variants, E1-V80L and E1-V80Q led to a reduction in specific infectivity (Figure 1B) and small plaque sizes on Vero cells (Figure 1C), suggesting that these variants may impact virion production. To rule out any effects of these substitutions on viral RNA synthesis we took advantage of a CHIKV reporter virus expressing luciferase under the control of the sub-genomic promoter such that luciferase will only be produced during active CHIKV genome replication (**Supplementary Figure 2A**). We transfected BHK-21 cells with each viral RNA and measured luciferase expression at 4, 6 and 8 hours post-transfection (**Supplementary Figure 2B**). We confirmed that the luciferase levels measured in the early time points (4 and 6 hours) are strictly associated to viral replication as infectious virus cannot be detected until 8 hours post-transfection (**Supplementary Figure 2B**). We used a non-replicative nsP4 active site polymerase mutant (GNN) to set the basal levels of noise. We found no differences in viral RNA replication between the different mutants (**Supplementary Figure 2B**) indicating that mutations E1-V80L and E1-V80Q are not playing a role in replication in BHK-21 cells and suggesting that these variants impair CHIKV virion production.

To confirm this, we performed viral growth curves at a multiplicity of infection (MOI) of 0.01 on both mammalian (BHK-21 and human foreskin fibroblasts (HFF-1)) and insect (C6/36 and *Ae. aegypti* (Aag2)) cells. We found that the E1-V80L and E1-V80Q substitutions significantly attenuated viral production in mammalian (Figure 1D) and mosquito cell lines (Figure 1E). Interestingly, E1-V80I and E1-V80A substitutions show increased viral production in HFF-1 cells, which is in line with what we have previously shown for E1-V80I in mouse embryonic fibroblasts and *in vivo* (Stapleford et al., 2014). All together these results show that altering residue E1-80, even with small changes in hydrophobicity, can modulate CHIKV virion production and infectivity *in vitro*.

### E1-V80L attenuation is conserved within alphaviruses

The valine residue at position 80 of the E1 glycoprotein is fully conserved within the alphavirus genus (Figure 1F). Thus, we asked whether we can attenuate other alphaviruses such as O’nyong’nyong virus (ONNV) and the distantly related alphavirus Sindbis virus (SINV) by introducing a leucine at position E1-80. Passages in BHK-21 cells demonstrated that, while SINV E1-V80L remained stable, the ONNV E1-V80L variant obtained a second-site mutation at position 65 (E1-T65S). Thus, we addressed the impact of E1-V80L substitution on the *in vitro* infectivity of SINV. We infected BHK-21 and C6/36 at a MOI of 0.01, and quantified infectious particles at 24 hours post infection. We observed attenuation in the infectivity of the SINV E1-V80L variant compared to the wildtype counterpart in both mammal and mosquito cell lines (Figure 1G). These data show that E1 position 80 is a key determinant of infectivity in not only CHIKV but in other alphaviruses as well.

### CHIKV E1-V80L and E1-V80Q reduce viral infection and dissemination in *Aedes aegypti* mosquitoes

Given the reduction of viral growth in mosquito cell lines, we hypothesized that these variants may impact CHIKV infection and dissemination in mosquitoes. We first infected *Ae. aegypti* mosquitoes with a high-dose artificial blood meal (10^6^ PFU/ml) of wildtype E1-V80, E1-V80A as an unbiased control, and E1-V80L and allowed infection to progress for 7 days. We found that although the E1-V80L variant is able to infect *Ae. aegypti* mosquitoes at similar levels compared to wild type (Figure 2A**, upper panel)**, there was a significant reduction in the amount of infectious virus in the mosquito bodies (Figure 2A**, lower panel)**. Moreover, viruses bearing the E1-V80L substitution showed impaired dissemination to legs and wings, with infectious particles detected in only 19 of the 44 infected mosquitoes (Figure 2A**, upper panel)**. Interestingly, we found that although E1-V80A had no impact on virus infectivity or dissemination (Figure 2A**, upper panel)**, there was a significant reduction in infectious virus in the legs and wings (Figure 2A**, lower panel)**.

**Figure 2.**
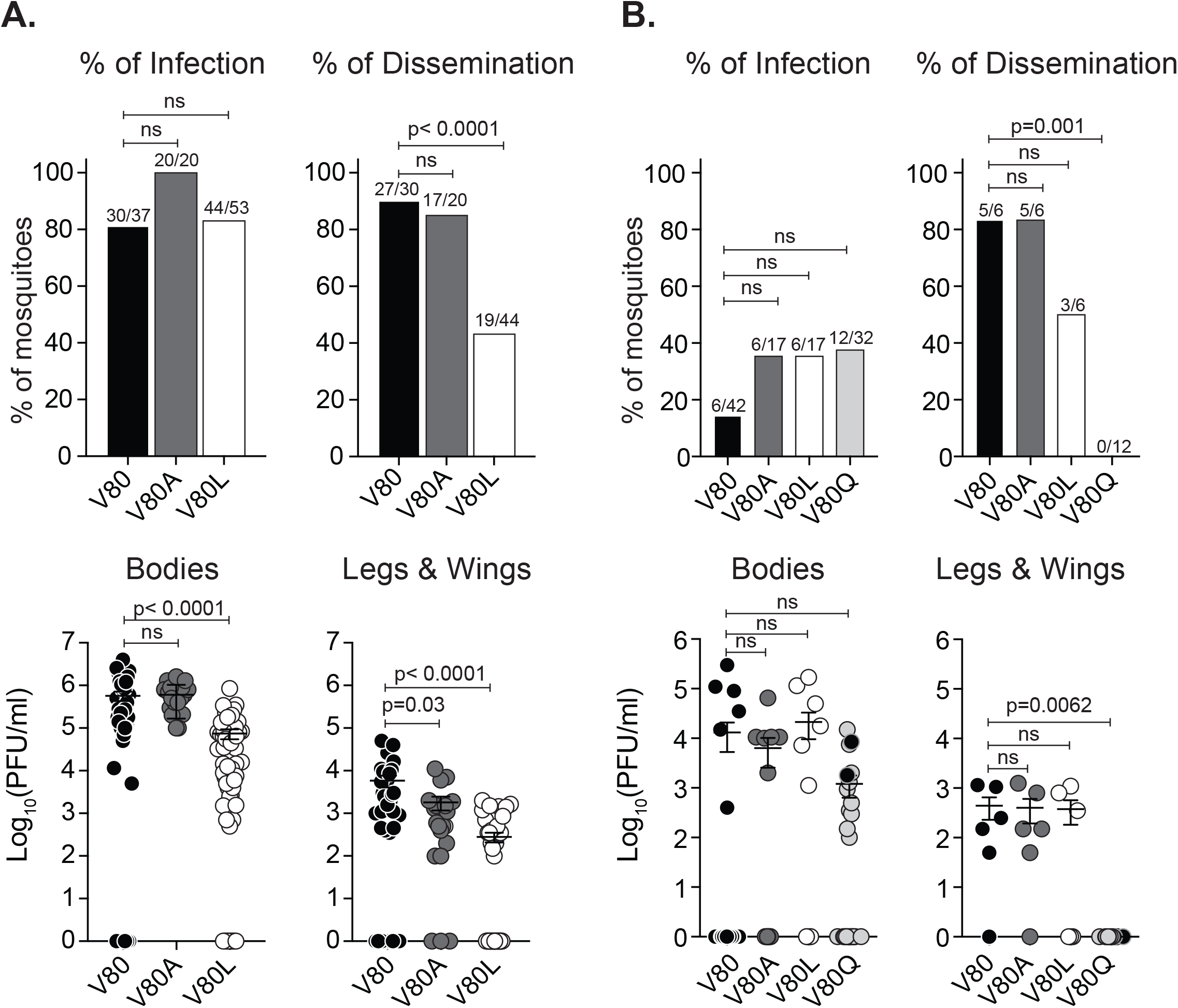
Chikungunya virus E1-V80L and E1-V80Q variants are attenuated and showed impaired dissemination in *Ae. aegypti* mosquitoes. 7 day old *Ae. aegypti* mosquitoes were infected with 10^6^ PFU/ml (A) or 10^4^ PFU/ml (B) of CHIKV E1-V80, V80A, V80L (A and B) and E1-V80Q (B) via artificial blood-meals. Viral titers of bodies and legs and wings were determined after 7 (A) or 14 (B) days post infection by plaque assay on Vero cells. Error bars represent mean and SEM, (N=17-53 mosquitoes). P values were determined by Kruskal-Wallis test. Percentage of infection or dissemination were determined as the number of mosquitoes with positive bodies or legs and wings over the total of engorged mosquitoes or positive bodies, respectively. P values were determined by Fisher’s exact test with Bonferroni correction. E1-V80Q reverted viruses are represented as black circles.

To address the impact of the variant E1-V80Q, for which we were unable to obtain high titer stocks, we infected mosquitoes with a low-dose of each virus (10^4^ PFU/ml) and allowed the infections to progress for 14 days. We found that at low doses, the variants had a trend towards higher infection rates compare to wildtype (Figure 2B**, upper panel)** and E1-V80Q yielded lower, although not statistically significant viral loads in the bodies of mosquitoes (Figure 2B**, lower panel).** Interestingly, based on the distinctive plaque size observed for E1-V80Q (Figure 1C) we found that viruses isolated from two mosquitoes reverted in plaque size (**Supplementary Figure 3A**). We Sanger sequenced the full viral structural region from these two mosquitoes and found that in addition to the E1-V80Q mutation, each individual virus acquired an additional second-site mutation in the E1 glycoprotein: E1-N20Y or E1-M88L (**Supplementary Figure 3A**). Both residues E1-20 and E1-88 are located in regions implicated in modulation of viral fusion (Voss et al., 2010; Zheng et al., 2011), suggesting that the attenuation observed for these mutants could be associated to a defect in fusion (**Supplementary Figure 3B and 3C)**. Importantly, of all the mosquitoes infected with E1-V80Q, including the ones with reversion in their plaque phenotype, none were able to disseminate into the legs and wings and this effect was also observed for E1-V80L although less pronounced (Figure 2B). Taken together, these studies indicate that position E1-80 is important for CHIKV infection and dissemination in *Ae. aegypti* mosquitoes and may function through impacting viral fusion.

### CHIKV E1-V80L and E1-V80Q are attenuated in mice and impair dissemination to secondary organs

In addition to reduced viral replication and impaired dissemination in mosquitoes we also observed that E1-V80L and E1-V80Q were impaired in mammalian cells and thus we asked whether these mutations attenuate CHIKV in mice. We infected adult C57BL/6J mice with 1000 PFU of E1-V80, V80A, V80L, and V80Q in the left footpad and addressed infectious viral loads and dissemination two days post infection which has been described as the peak of viremia of CHIKV for this model (Gardner et al., 2010) (Figure 3A). Interestingly, we found reduced levels of E1-V80L in the ipsilateral muscle (Figure 3B), a primary site of infection, as well as in the spleen and liver (Figure 3C), secondary sites of infection. Strikingly, we were unable to recover infectious E1-V80L particles from the heart and brain of infected mice indicating that this virus was significantly impaired in dissemination to specific organs (Figure 3C**, lower panel)**. On the other hand, we found that the E1-V80Q variant had reduced titers at the site of injection (Figure 3B) and we did not detect infectious virus in the serum or other secondary organs suggesting a strong restriction of this virus to the site of injection (Figure 3C).

**Figure 3.**
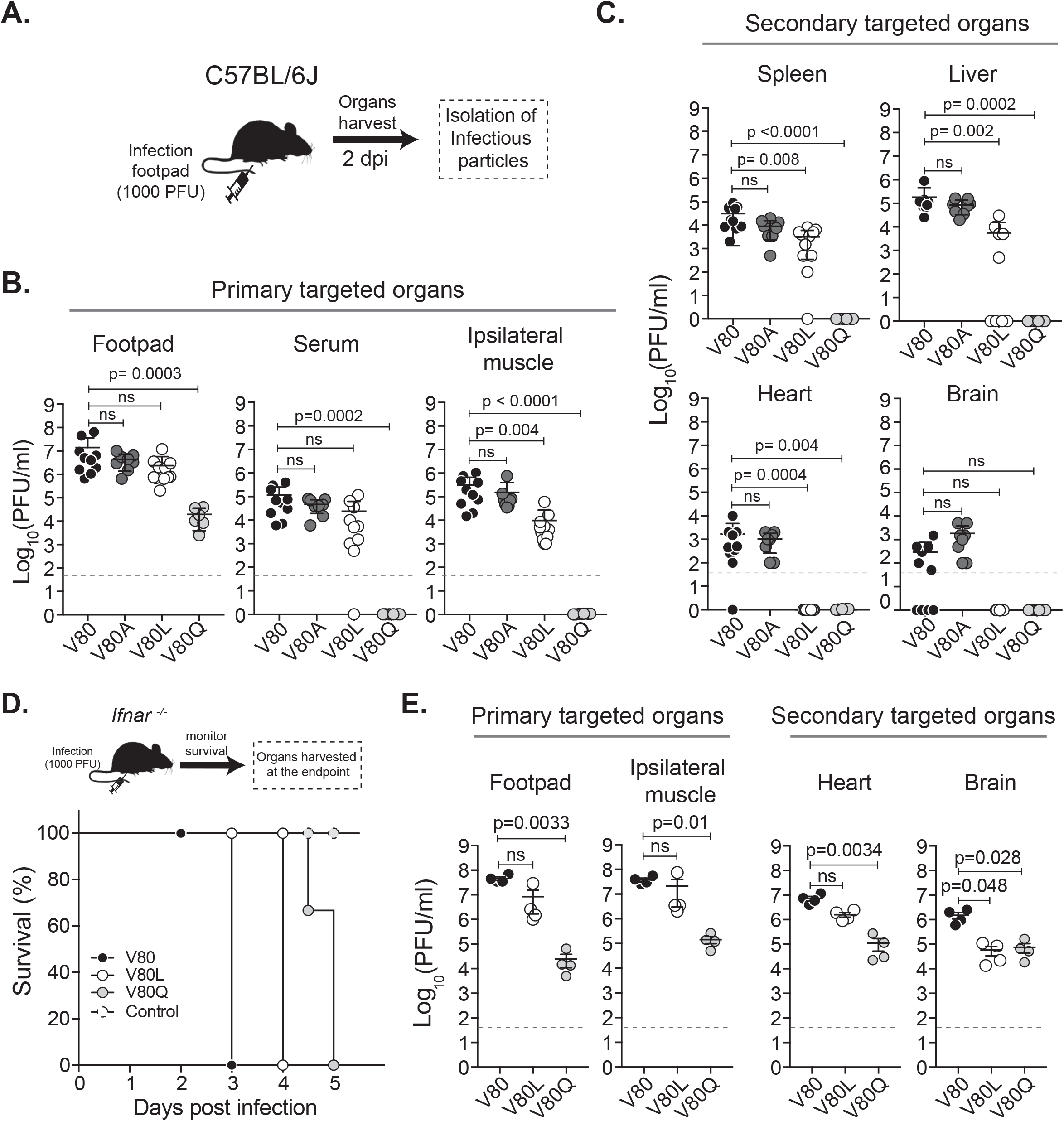
Chikungunya virus E1-V80L and E1-V80Q variants are attenuated and showed impaired dissemination in mice. (A) Schematic representation of the workflow. C57BL/6J mice were inoculated via footpad injection with 1000 PFU of E1-V80, V80A, V80L or V80Q variants. Two days post infection organs were harvested and viral loads were determined by plaque assay on Vero cells. Viral infectious particles were determined in the primary targeted organs (B) or secondary targeted organs (C). Results are expressed as PFU/ml of tissue homogenate. E1-V80 (N=10), E1-V80A (N=8), E1-V80L (N=11) and E1-V80Q (N=6). Error bars indicate SEM. All P values were determined by Kruskal-Wallis test. (D) Kaplan-Meier survival curves of C57BL/6J *Ifnar ^−/-^* mice inoculated via footpad injection with 1000 PFU of E1-V80 (black dots), E1-V80L (open dots), E1-V80Q (gray dots) or DMEM control (dashed dots) and monitored twice a day during 5 days (E1-V80L: p= 0.0009, E1-V80Q: p<0.0001, N=4; log-rank Mantel-Cox test). (E) Number of viral infectious particles was determined at the endpoint of the experiment by plaque assay on Vero cells. Results are expressed as PFU/ml of tissue homogenate. Error bars indicate SEM. P values were determined by Kruskal-Wallis test (N=4 per group). The plaque assay detection limit for these experiment is indicated as a dashed line.

One possible explanation for the restriction in dissemination, especially for the E1-V80Q variant is that the virus was not able to replicate in the injection site therefore not able to establish a systemic infection. To verify that E1-V80Q and E1-V80L viruses were infectious albeit attenuated, we took advantage of a susceptible mouse model lacking the interferon α/β receptor (*Ifnar^−/-^)* (Gardner et al., 2012). We infected *Ifnar^−/-^* mice with 1000 PFU of E1-V80, V80L, V80Q and a carrier control via footpad injection and monitored mice daily for signs of disease (**Supplemental Figure 4**). All mice succumbed to the infection compared to the uninfected control (Figure 3D); however, we observed an attenuation of E1-V80L and E1-V80Q through the delay in symptoms and death compared to wild type (**Supplementary Figure 4 and** Figure 3D). To examine the dissemination to the different organs, we analyzed the infectious viral loads in the primary and secondary target organs at the humane endpoints (Figure 3E). Notably, both E1-V80L and E1-V80Q were able to disseminate to all the tested organs, including heart and brain, but still showed attenuation compare to wildtype (Figure 3E). Taken together, these data show that E1-V80L and E1-V80Q variants are indeed infectious *in vivo* and that residue E1-80 plays a critical role in viral dissemination in both insects and mice.

### CHIKV residue E1-V80 impacts virion fusion dynamics and pH sensitivity

Given the proximity of residue E1-80 to the E1 glycoprotein fusion loop (Figure 1A) as well as the identification of second-site mutations (E1-V226A, E1-N20Y, and E1-M88L) in regions implicated in modulation of viral fusion (Voss et al., 2010; Zheng et al., 2011), we hypothesized that the viral attenuation and the impaired dissemination observed for the E1-V80 variants can be potentially associated with defects in viral entry.

To test this hypothesis, we generated ZsGreen reporter viruses of E1-V80, V80L, V80Q and V80A to visualize infected cells and preformed cellular binding and fusion assays (**Supplementary Figure 2A**). E1-V80Q variant was excluded from this study due to the inability to reach concentrated titers. We first addressed virion binding to mammalian cells by incubating BHK-21 cells on ice with gradient purified virions at a MOI of 0.1, and measuring the amount of bound virus after 5 and 60 minutes by RT-qPCR. We found that after 60 minutes E1-V80L showed reduced binding compared to wildtype (**Supplementary Figure 5**). Next, we performed fusion-from-without experiments on BHK-21 cells (Stapleford et al., 2014). In order to attempt to minimize the binding defect observed for E1-V80L we used a MOI of 10 for this assay. We found that E1-V80 and E1-V80A showed a threshold pH for fusion of ∼5.9, similar to what has been previously described for CHIKV (Hoornweg et al., 2016; Sanchez-San Martin et al., 2013; Stapleford et al., 2014). Remarkably, the fusion of E1-V80L with host cells was significantly reduced at multiple pH treatments compared to controls (Figure 4A), shifting the threshold pH for fusion to ∼5.5. As an alternative approach to look at pH dependence, we took advantage of the lysosomotropic agents ammonium chloride and bafilomycin A1 which deacidify the endosomal compartments and block viral fusion (Kielian et al., 1984; Sourisseau et al., 2007). We found that E1-V80L was more sensitive to each of these compounds compared to wild type confirming that this residue is able to sense and respond to pH changes (Figure 4B). Interestingly, we found that E1-V80A was sensitive to ammonium chloride treatment yet this may be through off target effects of the compound as we do not see this inhibition with bafilomycin A1, a selective inhibitor of endosome acidification. All together these data indicate that position E1-80 is determinant of viral fusion and that altering this residue can modulate viral fusion dynamics.

**Figure 4.**
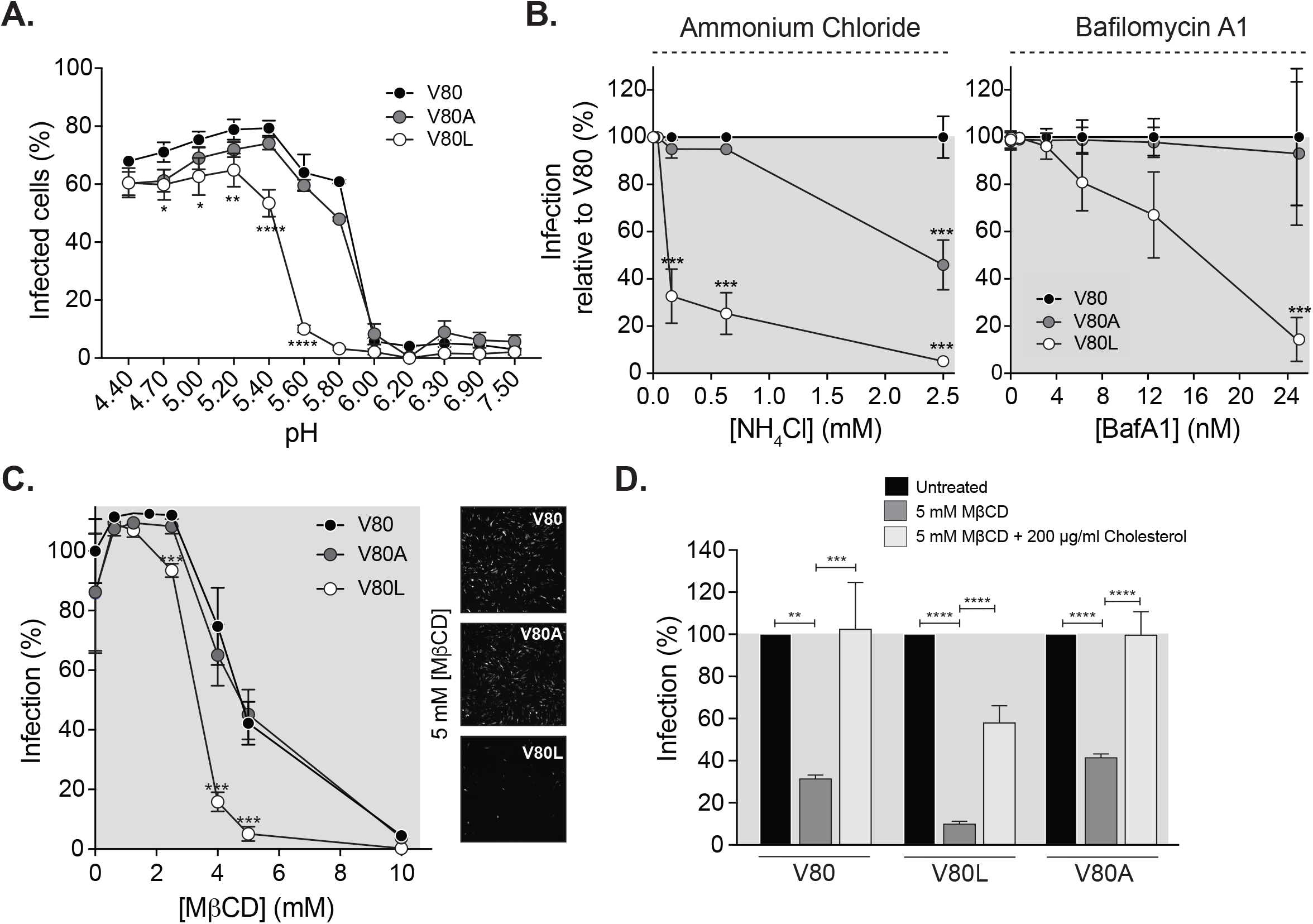
CHIKV residue E1-80 impacts virion fusion dynamics, pH sensitivity and cholesterol dependence. (A) CHIKV fusion from without. ZsGreen viruses were bound to BHK-21 cells at a MOI of 10 for one hour at 4 °C. After binding, cells were treated at the indicated pHs at 37 °C for 2 minutes. pH was neutralized, and the amount of ZsGreen expressing cells was determined after 16 hr post-infection. (B) Effect of lysosomotropic drugs on CHIKV mutant variants. BHK-21 cells were pre-treated for 3 hrs with increasing concentrations of ammonium chloride (left panel) or Bafilomycin A1 (right panel), and then infected with ZsGreen-viruses at a MOI of 1. The impact of each drug on viral infection was determined after 16 hr post infection. The mean and SEM are shown; N=3; (* p<0.05, ** p<0.01, *** p < 0.001). P values were determined by two-way ANOVA with Bonferroni post-hoc test. (C) Cholesterol depletion. BHK-21 cells were treated with increased concentrations of Methyl-β-cyclodextrin (MβCD) for one hr at 37 °C. Infections with ZsGreen-viruses were performed at a MOI of 1 for one hour, and complete media with 20 mM ammonium chloride was added to impair viral spreading. After 16 hr post-infection the amount of ZsGreen-expressing cells were determined. The mean and SEM are shown; N=4, (***p < 0.001). P values were determined by two-way ANOVA with Bonferroni post-hoc test. (D) Cholesterol repletion. BHK-21 cells were treated with 5mM MβCD for one 1hr at 37 °C. After treatment, cells were treated with media containing 200 μg/ml soluble cholesterol or media alone. Infections with ZsGreen-viruses were performed at a MOI of 1 for one hour, and complete media with 20 mM ammonium chloride was added to impair viral spreading. After 16 hr post-infection the amount of ZsGreen-expressing cells was determined. The mean and SEM are shown; N=6, (** p<0.01, *** p<0.001, **** p < 0.0001; One-way ANOVA) with Bonferroni multiple comparisons test.

### Residue E1-80 is involved in CHIKV E1 cholesterol dependence

It has been described for alphaviruses that during the pH-triggered E1 conformational transition, the fusion loop insertion and fusion is promoted by the presence of cholesterol in the target membranes (Guardado-Calvo et al., 2017; Kielian et al., 2000). Since we have shown that residue E1-80 impacts fusion dynamics, we next asked whether this residue may be sensitive to the cholesterol levels of the target membrane. To address this, we depleted cholesterol from BHK-21 cells with methyl beta-cyclodextrin (MβCD) (Hoornweg et al., 2016; Mahammad and Parmryd, 2015; Zidovetzki and Levitan, 2007), infected cholesterol-depleted cells with each virus variant to allow normal entry, added back media with 20 mM ammonium chloride to inhibit spread after infection and therefore measure the impact of cholesterol only during early stages in the viral infection. We found that while cholesterol depletion had no effect on the E1-V80A infectivity compared to wildtype, E1-V80L was significantly affected by this treatment (Figure 4C), indicating that this attenuated variant has increased cholesterol dependence. In order to confirm that the reduced infectivity observed for E1-V80L in the presence of MβCD was due to the effect of cholesterol, we replenished BHK-21 depleted cells with 200 μg/ml of cholesterol. We found that upon replenishment of membrane cholesterol, E1-V80L infectivity was restored (Figure 4D), indicating that residue E1-80 is involved in CHIKV cholesterol dependence.

### E1 residues 80 and 226 function together in CHIKV infectivity and dissemination

Position E1-226 has been shown to modulate the cholesterol requirement for CHIKV, SFV and SINV viral entry (Lu et al., 1999; Tsetsarkin et al., 2011; Vashishtha et al., 1998). Given the selection of the E1-V226A second-site variant during the E1-V80L adaptation in C6/36 cells (Table 1 **and Supplementary Figure 1B**), and that E1-V226A substitution in CHIKV IOL results in decreased cholesterol dependence for fusion (Hoornweg et al., 2016), we hypothesized that the acquisition of the E1-V226A mutation in CHIKV would rescue the E1-V80L cholesterol dependence, providing a functional link between these two residues. To test this hypothesis, we first introduced the E1-V226A mutation in our E1-V80L infectious clone, generated the double mutant virus (E1-V80L:V226A) and confirmed stability after three passages on BHK-21 cells. We found that the E1-V80L:V226A variant showed an increased plaque size compared to E1-V80L (Figure 5A), suggesting that this mutation rescued the impaired viral spreading observed for E1-V80L. We next addressed the capacity of E1-V80L:V226A to restore E1-V80L reduced infectivity in cholesterol-depleted cells. Indeed, we found that the E1-V80L:V226A variant reduced the cholesterol dependence observed for E1-V80L (Figure 5B), confirming that both residues E1-80 and 226 are involved in this process. Finally, we asked whether the E1-V226A substitution is able to rescue the E1-V80L *in vivo* attenuation and dissemination phenotype observed in mice. For that purpose, we infected C57BL/6J mice via footpad injection with ∼1000 PFU of E1-V80, V80L or V80L:V226A variants. We found that E1-V80L:V226A double mutant rescued the attenuated phenotype observed for E1-V80L in primary organs (Figure 5C). Interestingly, our results also indicate that the E1-V80L:V226A double mutant rescued the dissemination phenotype observed in the heart and the brain (Figure 5D), suggesting functional link between these two residues. Taken together, these results demonstrate that the highly conserved position 80 in the E1 glycoprotein is a key determinant in alphavirus infectivity and dissemination functioning in concert with E1 position 226 to modulate viral fusion and cholesterol dependence.

**Figure 5.**
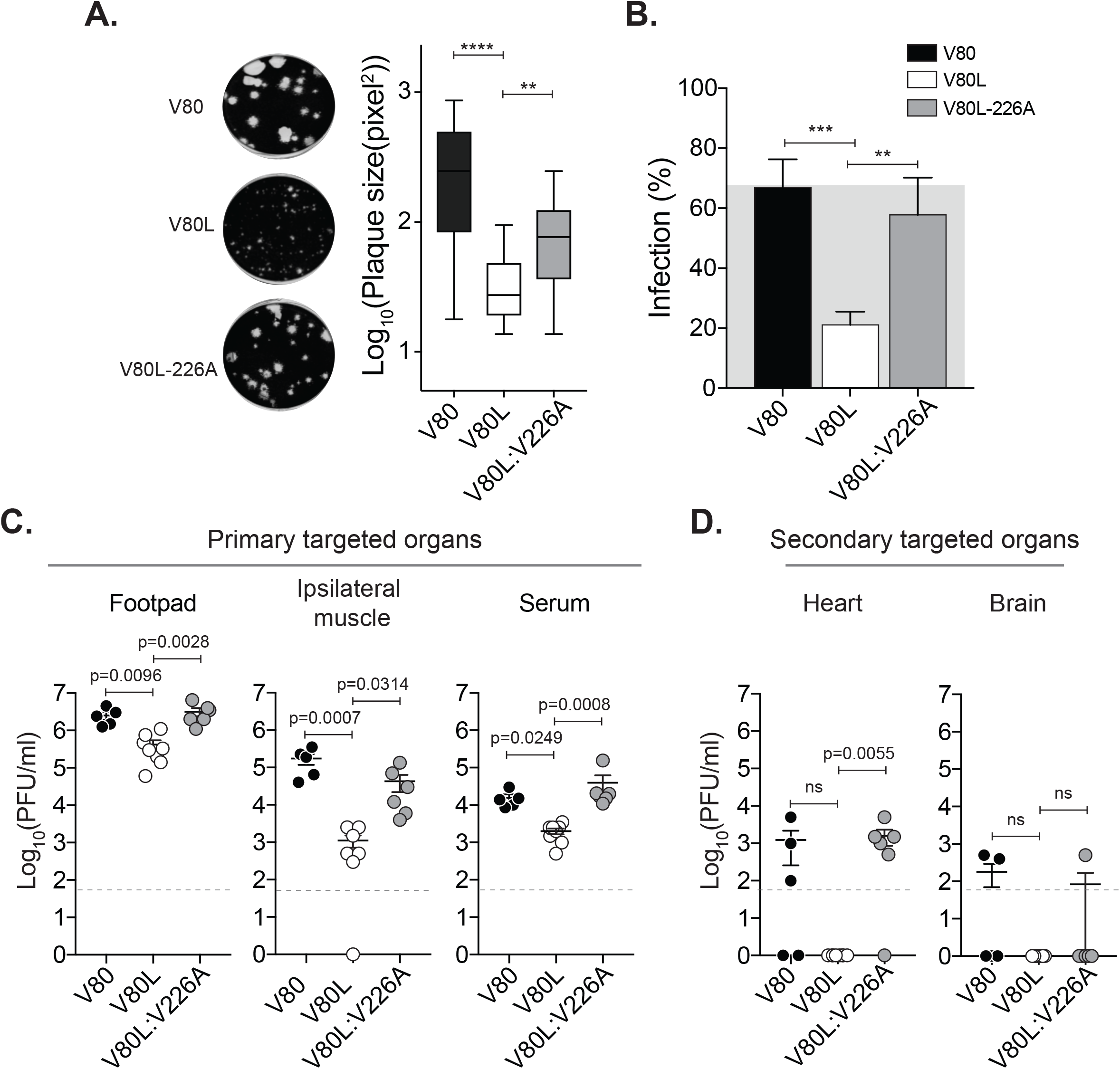
E1 residues 80 and 226 function together in CHIKV infectivity and dissemination. (A) Plaque size in Vero cells was quantified using ImageJ (**** p < 0.0001, ** p< 0.01, N > 40; Kruskal-Wallis test). (B) BHK-21 cells were treated with 5mM MβCD for one hr at 37 °C. Infections with ZsGreen-viruses were performed at a MOI of 1 for one hour, and complete media with 20 mM ammonium chloride was added to impair viral spreading. After 16 hr post-infection the amount of ZsGreen-expressing cells was determined. The mean and standard errors of the mean are shown; N=6, (*** p = 0.0006, ** p = 0.011; One-way ANOVA) with Bonferroni post-hoc test. (C-D) Mice were inoculated via footpad injection with 1000 PFU of V80, V80L or V80L:V226A variants. Two days post infection organs were harvested and viral loads were determined by plaque assay on Vero cells. Viral infectious particles were determined in the primary targeted organs (C) or secondary targeted organs (D). Results are expressed as PFU/ml of tissue homogenate. V80 (N=5), V80L (N=8), V80L:V226A (N=6). Error bars indicate SEM. All P values were determined by Kruskal-Wallis test.

## DISCUSSION

Arboviral diseases are undergoing a global expansion (Vignuzzi and Higgs, 2017; Wilder-Smith et al., 2017), pointing to the urgent need to study the fundamental aspects of arboviral biology to identify new therapeutic targets. We have previously developed an experimental approach that allowed us to identify position E1-80 as a potential hotspot of viral transmission and pathogenesis (Stapleford et al., 2014). In the present work, we performed a functional analysis of the highly conserved position 80 in E1 glycoprotein of CHIKV, by mutational tolerance studies and performing *in vitro* evolution in mammal and mosquito cell lines. We found that the E1-80 residue is a key determinant for viral infectivity and dissemination through the modulation of viral fusion and cholesterol dependence.

Position 80 in the E1 glycoprotein is located within the *β-strand c* at the tip of domain II, adjacent to the fusion loop, and in close proximity to a structurally conserved glycerophospholipid binding pocket (GPL) (Figure 6) (Guardado-Calvo et al., 2017; Voss et al., 2010). Our studies show that by changing this single amino acid we can regulate CHIKV infectivity by either increasing infectivity, as observed for E1-V80A and V80I in HFF-1 cells and pathogenesis, as observed for E1-V80I *in vivo* (Stapleford et al., 2014), or by attenuating viruses *in vitro* and *in vivo* as seen for E1-V80L and E1-V80Q. This ability to modulate viral infectivity provides powerful tools to study the fundamental mechanisms of arbovirus transmission and pathogenesis. In particular, the increased *in vitro* infectivity observed for the E1-V80A and V80I in HFF-1 cells, which have a competent innate immunity signaling pathway (Chen et al., 2017) is intriguing and suggests that this position can potentially be involved in regulatory processes other than fusion. Notably, this phenomenon of viral attenuation by position 80, is not exclusive to CHIKV, as we observed similar levels of attenuation in SINV, another member of the alphavirus genus. Such observations have strong implications in a generalized function of the E1-80 residue as an arboviral modulator of infectivity. Further research aimed towards defining analog residues in other arboviral type II glycoproteins would help to expand this concept.

**Figure 6.**
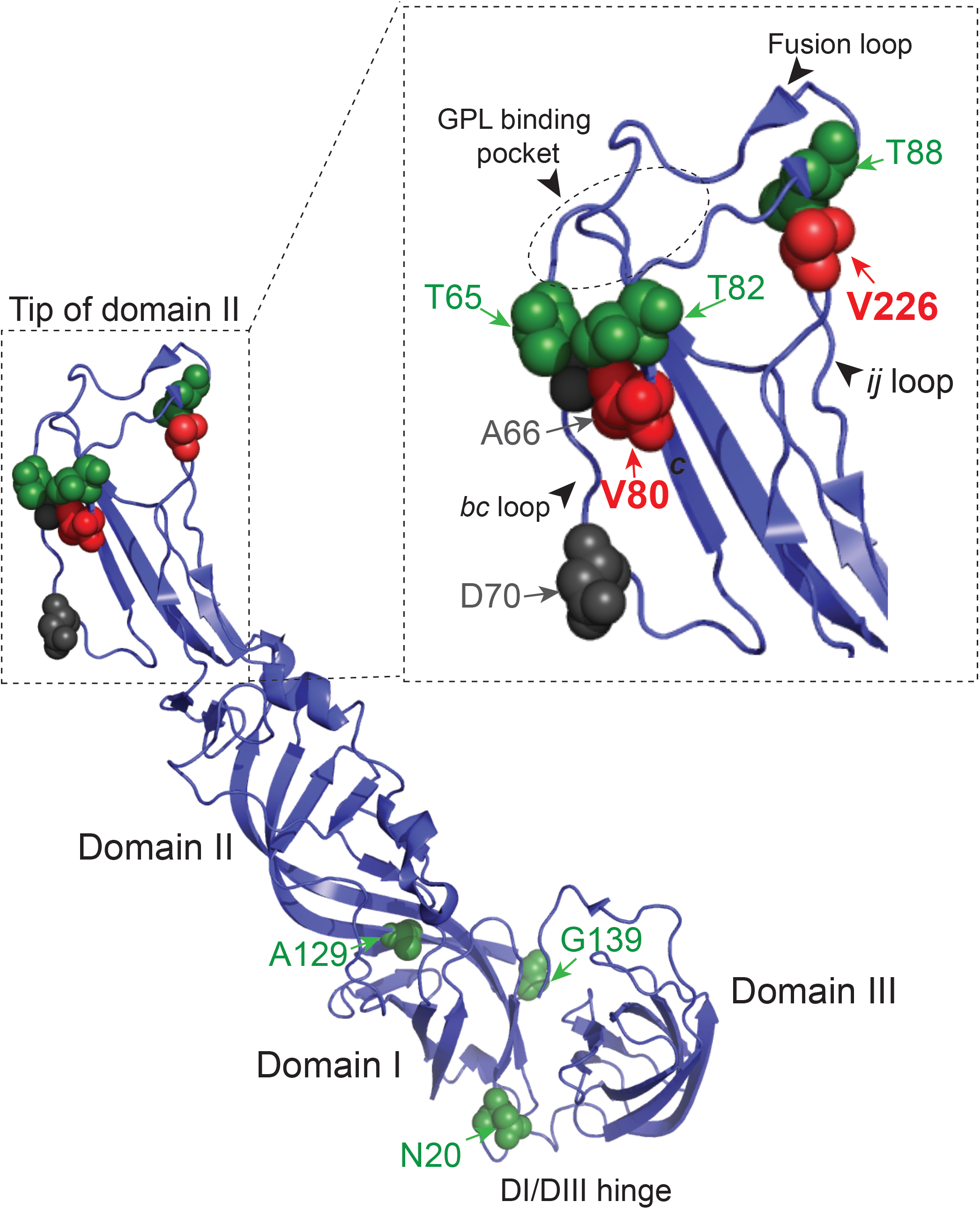
Summary of the identified second site mutations. Ribbon representation of the E1 CHIKV glycoprotein (PDBID: 3N42) indicating the mutated residues found in this study is shown. Second site mutations found in this studies are represented as green, black and red spheres. Detail of the tip of domain II (residues 53-108 and 218-235) is shown in the inset. In red are highlighted residues E1-80 and E1-226. Residues E1-66 and E1-70, corresponding to cholesterol dependence mutants, are highlighted in gray. Positions corresponding to the fusion loop, *ij loop*, *bc-loop, β-strand c* and the GPL binding pocket are indicated.

Our mutagenesis analysis showed host-specific constrains acting on E1-80, with the modified residues within the tip of domain II (E1-T65, A66, D70, T88, F95 and V226) or proximal to the hinge region between domain I and III (E1-N20, G139 and A129) (Figure 6), suggesting that each group of residues could be functionally linked. Notably, both the tip of domain II, as well as the hinge region between DI and DIII, are associated to membrane fusion (Kielian and Rey, 2006; Zheng et al., 2011), albeit through different mechanisms. While the tip of domain II is implicated with insertion of the fusion loop into the target membrane (Gibbons et al., 2003; Gibbons and Kielian, 2002) and interaction with lipids (Guardado-Calvo et al., 2017), the linker region between DI and DIII, has been associated to the stabilization of the post-fusion hairpin during the low-pH-dependent refolding (Zheng et al., 2011). In addition to coding changes, we also observed the emergence of synonymous mutations such us E1-A66A or E1-F95F, which could potentially be associated to RNA structural rewiring in response to the emergence of the new mutations. This, in addition with E1-V80A and E1-V80I increased infectivity leads us consider the possibility that RNA structure around position E1-80 may be impacting the CHIKV life cycle as well. Further work should be done in this regard, but we can hypothesize that there is an evolutionary trade-off between protein and RNA secondary structure constrains around position E1-V80 that might be impacting CHIKV evolution.

Nonetheless, our data demonstrates that E1-80 is an essential determinant of viral fusion by itself, and altering this residue modulates viral fusion dynamics. Thus, position E1-80 and the compensatory mutations that emerged during the *in vitro* passages can potentially be involved in a common mechanism regulating viral fusion. We speculate that compensatory mutations within the fusion pocket are associated with the fusion loop insertion and/or lipid binding, where the second-site mutations in the hinge region could be associated to a different mechanism, possibly favoring fusion by stabilizing the prefusion-post-fusion conformational transition. However, direct interactions between E1-E2 glycoproteins in the regulation of these processes can also provide a potential explanation for the observed phenotypes.

It has been widely described that the fusion loop insertion and fusion are promoted by the presence of cholesterol in the target membranes (Kielian et al., 2000). In this study, we demonstrated that position 80 of E1 glycoprotein modulates CHIKV cholesterol dependence in mammalian cell lines. In addition, passaging of E1-V80 variants on C6/36 drives the emergence of residues E1-226, E1-70 and E1-66 as second-site mutations (Figure 6 **and Supplementary Figure 1**), which were previously associated to cholesterol dependence of CHIKV infectivity (Tsetsarkin et al., 2011; Tsetsarkin et al., 2007). In particular, selection experiments on the attenuated E1-V80L and E1-V80Q variants in C6/36 cells strongly favor the emergence of E1-V226A variant. One potential explanation is that C6/36 cell lines, due to their reduced cholesterol levels (Hafer et al., 2009), are imposing a stronger selective pressure during *in vitro* evolution, resulting in a higher selective advantage of the E1-V226A mutation, whereby this modification compensates the reduced infectivity observed for E1-V80L. Indeed, our genetic studies support this hypothesis, given that the incorporation of E1-V226A into the E1-V80L infectious clone rescued viral spreading, cholesterol dependence and infectivity *in vivo* (Figure 5). In this study, we have found a new region implicated in modulation of cholesterol dependence at the tip of domain II, which involves position 80. We speculate, that the *ij* loop (containing residue 226), the *β-strand c* (containing residue 80) and the *bc loop* (containing residues 66 and 70) (Figure 6) work concerted to accommodate cholesterol within the tip of domain II during the fusion process. Recent structural studies have identified a conserved GPL binding pocket involving residues from the *bc loop*, fusion loop and *ij loop* in E1 glycoprotein essential for fusion loop insertion (Guardado-Calvo et al., 2017). In addition, the authors propose that additional small head group lipids (such as cholesterol) are required to create space between the bulky head group to allow fusion loop insertion. Structural studied will be need to address the role of position 80 in cholesterol dependence which could be implicated with direct E1-80 mediated interactions, or through modulation of the loops dynamic during the conformational transition and/or insertion in the target membrane.

Finally, we showed that in adult C57BL/6, both the E1-V80L and V80Q variants had impaired dissemination to secondary organs compared to wildtype CHIKV (Figure 3); however, this impairment was restored in *Ifnar^−/-^* mice. It has been described that decreasing *de novo* cholesterol and fatty acid biosynthesis is a physiological response to viral infection via the IFNAR signaling pathway (York et al., 2015). This suggests, that the impaired dissemination observed in wildtype mice could be associated to reduction of available cholesterol levels in the membrane during CHIKV infection, which resulted in impaired E1-V80L and E1-V80Q dissemination to target organs. Although the increased infectivity and dissemination observed in *Ifnar^−/-^* mice could be associated to many other regulatory processes, such as host specific innate and adaptive antiviral immunity (Nair et al., 2017), it will be interesting to study the impact of cholesterol levels on the E1-V80L and E1-V80Q variants in *Ifnar^−/-^*mouse models. Similar associations can be modulating the impaired dissemination observed in *Ae. aegypti*, suggesting that cholesterol levels in the midgut could be responsible for determining the degree of dissemination of these variants. However, more studies are required to test these hypotheses.

In summary, we show that the E1 residue 80 is a key determinant in alphavirus infectivity and dissemination through modulation of viral fusion and cholesterol dependence, and that it is functionally linked with other residues to orchestrate viral fusion, cholesterol dependence, infectivity and dissemination. In addition, these data provide evidence towards the potential use of the tip of domain II as a druggable target, gearing towards the development of broad spectrum of alphavirus antivirals and opening a new avenue towards the use of mutational tolerance approaches as a way to identify stable substitutions guiding the development of attenuated vaccines.

## Acknowledgements

We would like to thank all members of the Stapleford lab for helpful comments on this manuscript. We also thank Meike Dittmann, Maren de Vries and Aaron Briley for help with the CX7 CellInsight high-content microscope and analysis, and Ken Cadwell, Kamal Khanna and Stephen Yeung who kindly provided essential reagents. M.G.N was supported by a Jan Vilcek/David Goldfarb Fellowship from the New York University Department of Microbiology, New York University and M.R. is supported in part by the Public Health Service Institutional Research Training Aware T32 AI007180.

## Author Contributions

M.G.N. and K.A.S. designed experiments, M.G.N., B.A.R.R., M.R. and K.A.S. performed experiments, M.G.N. and K.A.S. analyzed the data, M.G.N. and K.A.S. wrote the paper.

## STAR Methods

### Cells and viruses

Mammalian cell lines were maintained at 37°C in 5% CO_2_. Baby hamster kidney (BHK-21, ATCC CCL-10) and Human foreskin fibroblast (HFF-1, ATCC SCRC-1041) cells were grown in Dulbecco’s modified Eagle’s medium (DMEM; Corning) supplemented with 10% fetal bovine serum (FBS; Atlanta biologicals), 1% nonessential amino-acids (NEAA, Gibco) and 1% penicillin-streptomycin (P/S, Gibco). Vero cells (ATCC CCL-81) were grown in DMEM with 10% newborn calf serum (NBCS; Gibco) and 1% P/S. Mosquitoes cell lines were maintained at 28°C in 5% CO_2_. *Ae. aegypti* cells (Aag-2 obtained from Dr. Paul Turner, Yale University) were grown in DMEM supplemented with 10% FBS and 10% NEAA. *Ae. albopictus* cells (C6/36, ATCC CRL-1660) were maintained in L15 medium (Corning) supplemented with 10% FBS, 1% tryptose phosphate broth, 1% NEAA, and 1% P/S. All the cell lines were confirmed free of mycoplasma.

Wildtype chikungunya virus (CHIKV) was rescued from the CHIKV strain 06-049 (AM258994) infectious clone as previously described (Coffey and Vignuzzi, 2011). To generate the CHIKV E1 glycoprotein position 80 variants, each amino acid substitution was first introduced into a cloning vector by site-directed mutagenesis using Phusion DNA polymerase (Thermo-Scientific). Primer sequences are provided in Supplemental Table 1. Each cloning plasmid was Sanger sequenced (Genewiz) from common 5’ XhoI and 3’ NotI restriction enzyme sites to confirm each mutation and the absence of second-site variants. The XhoI/NotI fragment of each cloning vector was subcloned into the CHIKV full-length infectious clone by standard molecular biology techniques and subsequently sequencing to confirm each mutation. A CHIKV La Reunion infectious clone expressing ZsGreen and Firefly luciferase was constructed by standard molecular biology techniques. First, an AvrII restriction enzyme site was inserted 5’ of the subgenomic promoter by site-directed mutagenesis using the primers (Forward 5’ CACTAATCAGCTACACCTAGGATGGAGTTCATCCC 3’ and Reverse 5’ GGGATGAACTCCATCCTAGGTGTAGCTGATTAGTG 3’). The CHIKV subgenomic promoter was then amplified by PCR (Forward 5’ CCTAGGCCATGGCCACCTTTGCAAG 3’ and Reverse 5’ ACTAGTTGTAGCTGATTAGTGTTTAG 3’) and subcloned into the AvrII site to generated a CHIKV infectious clone containing two subgenomic promoters. Finally, the ZsGreen and Firefly luciferase cassettes were amplified by PCR (Primers in Supplemental Table 1) from CHIKV infectious clones obtained from Andres Merits (University of Tartu) and subcloned into the AvrII restriction enzyme site. The complete cassette and subgenomic regions were sequenced to ensure there were no second-site mutations. ZsGreen and Firefly reporter constructs containing individual E1 mutations were constructed by subcloning the XhoI/NotI fragment from the unmarked infectious clones into similar sites in the ZsGreen and Firefly luciferase backgrounds. The Sindbis virus (SINV) and O’nyongn’yong virus (ONNV) infectious clones were obtained from Dr. Benjamin tenOever (Mt. Sinai Ichan School of Medicine) and Andres Merits respectively. E1-V80 glycoprotein variants were generated as described above.

To generate infectious virus, 10 μg of each CHIKV plasmid was linearized overnight with NotI (Thermo-Scientific), phenol:chloroform extractioned, ethanol precipitated, and resuspended in nuclease-free water at 1 μg/μl. *In vitro* transcribed viral RNAs were produced using the SP6 mMESSAGE mMACHINE kit (Ambion) following the manufacturer’s instructions. After RNA synthesis, samples were DNase treated and RNAs were purified by phenol:chloroform extraction, ethanol precipitation, and resuspended at a concentration of 1 μg/μl. All RNAs were stored at −80 °C until electroporation. SINV and ONNV infectious clones were digested with XhoI and PmeI respectively and prepared as describe above. Infectious virus was produced by electroporating BHK or C6/36 cells with *in vitro* transcribed viral RNAs. Briefly, cells were trypsinized, washed twice with ice-cold DPBS, and resuspended at 1×10^7^ cells/ml in PBS. 390 μl of cells were mixed with 10 μg of *in vitro* transcribed RNA and added to a 2 mm electroporation cuvette (BioRad). BHK-21 cells were electroporated with 1 pulse at 1.2 kV, 25 μF, with infinite resistance and C6/36 cells were electroporated with 1 pulse at 250 V, 550 μF, with a resistance of 25Ω. Cells were allowed to recover for 10 min at room temperature, transferred into 6 ml of warm DMEM, and placed in a T25 flask at 37 °C for 72 h. Virus was harvested, clarified at 1,200 × g for 5 min, and 3 ml of the clarified supernatants were used to infect T75 flask to generate working stocks (Passage 1). Viral titers were determined by plaque assay. In brief, 10-fold dilutions of each virus in DMEM were added to a monolayer of Vero cells for 1 hour at 37°C. Following incubation, cells were overlaid with 0.8% agarose in DMEM containing 2% NCBS and incubated at 37°C for 72 hours. The cells were fixed with 4% formalin, the agarose plug removed, and plaques visualized by crystal violet staining. Each mutant virus was generated in either BHK-21 or C6/36 cells three independent times. Each virus was passaged three times on each cell type it was generated in and viral RNA was extracted by TRIzol reagent (Invitrogen). Viral RNA was used directly for cDNA synthesis using the Maxima H minus-strand kit (Thermo). CHIKV cDNA was then used to generate two PCR amplicons (Fragment 1: spanning from 7203 to 10383 bp, Fragment 2: spanning from 9792 to 11541 bp), each fragment was purified using the Nucleospin PCR and Gel Extraction kit (Macherey-Nagel), and was Sanger sequenced (Genewiz) to address variant genetic stability and second-site mutations (**Supplementary Figure 1A**).

### Virus RNA replication and luciferase assay

BHK-21 cells (20,000 cells/well) were transfected with 90 ng of each viral RNA via Lipofectamine 2000 reagent (Invitrogen) following the manufacturer’s instructions. At each time point, the virus containing supernatant was removed, and 100 μl of Steady-Glo firefly luciferase reagent (Thermo) was added directly to the wells. Luminescence was quantified using the Perkin Elmer EnVision^TM^ plate reader under the US LUM settings. The virus containing supernatant was then placed on naïve BHK-21 cells for 1 hour to address virion production. After incubation, supernatant was removed, cells washed two times with PBS, and 100 μl of complete media was added to each well. Luminescence was quantified after 24 hours as described above.

### Viral growth curves and extracellular RNA levels

Viral growth was determined in BHK-21, C6/36, and Aag2 cells by infecting each cell type at an MOI of 0.01 in serum-free media at 37°C for 1 hour. Cells were washed extensively with PBS, complete media added, and virus containing supernatants were harvested at time 0, 4, 8, 12, 24, and 48 hours post infection. Viral titers were quantified by plaque assay as described above. For SINV, viral titers at 24 hours post infection were quantified by TCID_50_. In brief, Vero cells were seeded into 96-well plates (10,000 cells/well). 10-fold dilutions of each virus were made in DMEM and 100 μl of virus was added to each well. Virus-cell mix were incubated at 37 °C with 5% CO_2_ for 7 days. Following incubation, cells were fixed with 4% formalin, and visualized with crystal violet. In addition, viral RNA extractions were performed using TRIzol^TM^ Reagent (Invitrogen^TM^), and number of viral genomes were quantified by RT-qPCR using Taqman^®^ (Applied Biosystems^TM^) (Supplementary table 1).

### Plaque size quantification

Plaque assays were performed in 6 well plates and plaques were photographed using Chemidoc touch Imagin System (Biorad) at a fixed distance. Plaque sizes were measured in pixels using ImageJ software.

### Mosquito infections and harvests

*Aedes aegypti* mosquitoes (Poza Rica, Mexico, P20, obtained from Dr. Gregory Ebel, Colorado State University) were infected via artificial blood meals with high or low viral loads (10^6^ or 10^4^ PFU/ml, respectively). Briefly, viruses were mixed 1:2 with pre-washed rabbit whole blood (BioIVT) supplemented with 5 mM ATP. Female mosquitoes were allowed to feed on 37 °C blood meals through an artificial membrane for 60 to 120 minutes. Engorged females were identified, sorted and incubated at 28 °C with 10% sucrose ad libitum for 7 day or 14 days for high and low viral loads infections, respectively. After incubations, legs and wings were removed, and placed in 2 ml round bottom tubes with 500 μl of PBS containing one 5 mm stainless steel beads (QIAGEN), homogenized with TissueLyser II (QIAGEN), and debris was pulled down at 1200 rpm for 10 minutes. Viral titers in sample homogenates were determined by plaque assay using Vero cells.

### Mouse infections

6-9 week old male and female C57BL/6J and 6 week old *Ifnar^−/-^* mice were bred and grown in house. C57BL/6J and *Ifnar^−/-^*mice were anesthetized by inhalation of isoflurane (Henry Schein Animal Health), and infected via footpad injection with 10^3^ PFU of each mutant variant in 50 μl of DMEM. Mock-infected animals received DMEM alone. C57BL/6J mice were euthanized 2 days post infection by CO_2_ inhalation. *Ifnar^−/-^* were monitored for clinical signs of diseases twice a day and weighed at 24 hours intervals. After reaching the humane endpoint all the mice were euthanized by CO_2_ inhalation. Blood was collected by cardiac puncture. Organs including footpad, muscle, liver, heart, brain and spleen were collected in 500 μl of PBS containing one or two 5 mm stainless steel beads (QIAGEN), homogenized with TissueLyser II (QIAGEN), and debris was pulled down at 8,800 rpm for 10 minutes. Viral titers in tissue homogenates were determined by plaque assay using Vero cells. Animal experiments were performed in accordance with all NYU School of Medicine Institutional Animal Care and Use Committee guidelines (IACUC). All mouse studies were performed using biosafety level 3 conditions.

### Virus Binding assays

All ZsGreen viruses were first purified over a 20% sucrose cushion by ultracentrifugation at 25,000 rpm for 2 hours and resuspended in virus infection media (DMEM containing 0.2% BSA, 1 mM HEPES pH 7.4 and 1% P/S). For binding assays, BHK cells were preincubated with media containing 20 mM ammonium chloride for 2 hours at 4°C. Viruses were diluted in binding buffer (RPMI, 0.2% BSA, 10 mM HEPEs, 20 mM ammonium chloride) and incubated with BHK at a MOI of 0.1 for 5 or 60 mins at 4 °C. After incubation, the cells were washed extensively with ice-cold PBS, and RNA was extracted with Trizol reagent as described previously. CHIKV RNA was quantified by RT-qPCR as described above.

### Virus fusion from without assays

BHK cells (35,000 cells/well) were incubated with binding buffer (RPMI, 0.2% BSA, 10 mM HEPES, 20 mM ammonium chloride) for 1 hr at 4°C. Then, ZsGreen purified viruses were added at a MOI of 10 for 1 hr at 4°C. Unbound virus was removed and viral fusion was induced by adding fusion buffer for 2 min at 37°C (RPMI, 0.2% BSA, 30 mM succinic acid, 10 mM HEPES, adjusted to each pH described and prewarmed at 37°C). After incubation, the pH curve was removed and replaced with complete media with 20 mM ammonium chloride. After incubation at 37°C for 18 hrs, cells were fixed with 1% paraformaldehyde (PFA). After fixation, cells were permealized with 0.25% triton-100, stained with DAPI, and quantified on a CellInsight CX7 High-content microscope (Thermo) using a cut-off for three standard deviations from negative to be scored as an infected cell.

### Lysosomotropic drug sensitivity assays

BHK-21 cells (30,000 cells/well) were seeded in 96 well plates, 24 hours before treatment. Cells were then pretreated for 3 hours in serum free media containing increasing concentrations of ammonium chloride or balfilomycin A1 (Sigma). Following incubation, cells were infected with each ZsGreen virus at an MOI of 1 in the presence of each compound for 1 hour at 37 °C. Cells were washed extensively, complete media containing each compound was added, and cells were incubated at 37°C for 16 hours. In addition, control cells were treated with equivalent dilutions of the drug with no virus, and virus with no drug. After incubation, cells were fixed with 4% PFA and ZsGreen positive cells were quantified as described above.

### Cholesterol depletion and repletion experiments

BHK-21 cells were pretreated for 1 hour with increasing concentrations of methyl-beta-cyclodextrin (MβCD). Following incubation, cells were washed once with PBS and infected with each ZsGreen virus at an MOI of 1 for 1 hour at 37°C. The cells were then washed extensively, complete media containing 20 mM ammonium chloride was added, and cells were incubated at 37°C for 18 hours. After incubation, cells were fixed, stained, and ZsGreen positive cells were quantified as described above. For Cholesterol repletion experiments, cholesterol depleted cells were treated for 1 hour with 200 μg/ml of water-soluble cholesterol (Sigma), after incubation cells were infected and treated as described above.

### Sequence alignments and LOGOS

Sequence alignments were performed using the MUSCLE software with default parameters (Edgar, 2004). We generated sequence alignments of 13 representative members of the alphavirus genus: Chikungunya virus IOL (CHIKV, ACS29297.1), O’nyong’nyong virus (ONNV, NP_740711.1), Sindbis virus (SINV, NP_740677.1), Semliki forest virus (SFV, P03315), Sagiyama virus (SAGV, Q9JGK8), Venezuelan equine encephalitis virus (strain Trinidad donkey (VEEV, P09592), Aura virus (AURAV, Q86925), Eastern equine encephalitis virus (EEEV, P08768), Mayaro virus (strain Brazil) (MAYAB, Q8QZ72), Ross river virus (RRVN, P13890), Middelburg virus (MIDDV, Q80S27), Barmah forest virus (BFV, P89946) and Western equine encephalitis virus (WEEV, P13897). Sequence logos describing the conservation between residues 62 and 99 of E1 glycoprotein were generated with a sequence alignment of 13 representative members of the alphavirus genus using WebLogo software (Crooks et al., 2004). The x-axis indicates the position within the alignment and the y-axis indicates the information content. The height of each stack indicates conservation and the height of each letter is proportional to the frequency of that residue in that position in the alignment.

### Protein representations

All protein representations were performed using the program Pymol (https://pymol.org/).

### Statistical analyses

Statistics were performed using GraphPad Prism and R-studio software. P-values > 0.05 were considered non-significant (ns).

## REFERENCES

Brault, A.C., Powers, A.M., Ortiz, D., Estrada-Franco, J.G., Navarro-Lopez, R., and Weaver, S.C. (2004). Venezuelan equine encephalitis emergence: enhanced vector infection from a single amino acid substitution in the envelope glycoprotein. Proc Natl Acad Sci U S A 101, 11344–11349.

Chen, D., Long, M., Xiao, B., Xiong, Y., Chen, H., Chen, Y., Kuang, Z., Li, M., Wu, Y., Rock, D.L., et al. (2017). Transcriptomic profiles of human foreskin fibroblast cells in response to orf virus. Oncotarget 8, 58668–58685.

Coffey, L.L., and Vignuzzi, M. (2011). Host alternation of chikungunya virus increases fitness while restricting population diversity and adaptability to novel selective pressures. J Virol 85, 1025–1035.

Crooks, G.E., Hon, G., Chandonia, J.M., and Brenner, S.E. (2004). WebLogo: a sequence logo generator. Genome Res 14, 1188–1190.

Ebel, G.D., Carricaburu, J., Young, D., Bernard, K.A., and Kramer, L.D. (2004). Genetic and phenotypic variation of West Nile virus in New York, 2000-2003. Am J Trop Med Hyg 71, 493–500.

Gardner, C.L., Burke, C.W., Higgs, S.T., Klimstra, W.B., and Ryman, K.D. (2012). Interferon-alpha/beta deficiency greatly exacerbates arthritogenic disease in mice infected with wild-type chikungunya virus but not with the cell culture-adapted live-attenuated 181/25 vaccine candidate. Virology 425, 103–112.

Gardner, J., Anraku, I., Le, T.T., Larcher, T., Major, L., Roques, P., Schroder, W.A., Higgs, S., and Suhrbier, A. (2010). Chikungunya virus arthritis in adult wild-type mice. J Virol 84, 8021–8032.

Gibbons, D.L., Erk, I., Reilly, B., Navaza, J., Kielian, M., Rey, F.A., and Lepault, J. (2003). Visualization of the target-membrane-inserted fusion protein of Semliki Forest virus by combined electron microscopy and crystallography. Cell 114, 573–583.

Gibbons, D.L., and Kielian, M. (2002). Molecular dissection of the Semliki Forest virus homotrimer reveals two functionally distinct regions of the fusion protein. J Virol 76, 1194–1205.

Gould, E., Pettersson, J., Higgs, S., Charrel, R., and de Lamballerie, X. (2017). Emerging arboviruses: Why today? One Health 4, 1–13.

Greene, I.P., Paessler, S., Austgen, L., Anishchenko, M., Brault, A.C., Bowen, R.A., and Weaver, S.C. (2005). Envelope glycoprotein mutations mediate equine amplification and virulence of epizootic venezuelan equine encephalitis virus. J Virol 79, 9128–9133.

Guardado-Calvo, P., Atkovska, K., Jeffers, S.A., Grau, N., Backovic, M., Perez-Vargas, J., de Boer, S.M., Tortorici, M.A., Pehau-Arnaudet, G., Lepault, J., et al. (2017). A glycerophospholipid-specific pocket in the RVFV class II fusion protein drives target membrane insertion. Science 358, 663–667.

Hafer, A., Whittlesey, R., Brown, D.T., and Hernandez, R. (2009). Differential incorporation of cholesterol by Sindbis virus grown in mammalian or insect cells. J Virol 83, 9113–9121.

Higgs, S., and Vanlandingham, D. (2015). Chikungunya virus and its mosquito vectors. Vector Borne Zoonotic Dis 15, 231–240.

Hoornweg, T.E., van Duijl-Richter, M.K.S., Ayala Nunez, N.V., Albulescu, I.C., van Hemert, M.J., and Smit, J.M. (2016). Dynamics of Chikungunya Virus Cell Entry Unraveled by Single-Virus Tracking in Living Cells. J Virol 90, 4745–4756.

Kielian, M., Chatterjee, P.K., Gibbons, D.L., and Lu, Y.E. (2000). Specific roles for lipids in virus fusion and exit. Examples from the alphaviruses. Subcell Biochem 34, 409–455.

Kielian, M., and Rey, F.A. (2006). Virus membrane-fusion proteins: more than one way to make a hairpin. Nat Rev Microbiol 4, 67–76.

Kielian, M.C., Keranen, S., Kaariainen, L., and Helenius, A. (1984). Membrane fusion mutants of Semliki Forest virus. J Cell Biol 98, 139–145.

Labeaud, A.D., Bashir, F., and King, C.H. (2011). Measuring the burden of arboviral diseases: the spectrum of morbidity and mortality from four prevalent infections. Popul Health Metr 9, 1.

Lu, Y.E., Cassese, T., and Kielian, M. (1999). The cholesterol requirement for sindbis virus entry and exit and characterization of a spike protein region involved in cholesterol dependence. J Virol 73, 4272–4278.

Mahammad, S., and Parmryd, I. (2015). Cholesterol depletion using methyl-beta-cyclodextrin. Methods Mol Biol 1232, 91–102.

Moudy, R.M., Meola, M.A., Morin, L.L., Ebel, G.D., and Kramer, L.D. (2007). A newly emergent genotype of West Nile virus is transmitted earlier and more efficiently by Culex mosquitoes. Am J Trop Med Hyg 77, 365–370.

Nair, S., Poddar, S., Shimak, R.M., and Diamond, M.S. (2017). Interferon regulatory factor-1 (IRF-1) protects against chikungunya virus induced immunopathology by restricting infection in muscle cells. J Virol.

Pesko, K.N., and Ebel, G.D. (2012). West Nile virus population genetics and evolution. Infect Genet Evol 12, 181–190.

Powers, A.M. (2018). Vaccine and Therapeutic Options To Control Chikungunya Virus. Clin Microbiol Rev 31.

Rey, F.A., and Lok, S.M. (2018). Common Features of Enveloped Viruses and Implications for Immunogen Design for Next-Generation Vaccines. Cell 172, 1319–1334.

Rodriguez-Morales, A.J., Villamil-Gomez, W.E., and Franco-Paredes, C. (2016). The arboviral burden of disease caused by co-circulation and co-infection of dengue, chikungunya and Zika in the Americas. Travel Med Infect Dis 14, 177–179.

Sanchez-San Martin, C., Nanda, S., Zheng, Y., Fields, W., and Kielian, M. (2013). Cross-inhibition of chikungunya virus fusion and infection by alphavirus E1 domain III proteins. J Virol 87, 7680–7687.

Schuffenecker, I., Iteman, I., Michault, A., Murri, S., Frangeul, L., Vaney, M.C., Lavenir, R., Pardigon, N., Reynes, J.M., Pettinelli, F., et al. (2006). Genome microevolution of chikungunya viruses causing the Indian Ocean outbreak. PLoS Med 3, e263.

Silva, L.A., and Dermody, T.S. (2017). Chikungunya virus: epidemiology, replication, disease mechanisms, and prospective intervention strategies. J Clin Invest 127, 737–749.

Sourisseau, M., Schilte, C., Casartelli, N., Trouillet, C., Guivel-Benhassine, F., Rudnicka, D., Sol-Foulon, N., Le Roux, K., Prevost, M.C., Fsihi, H., et al. (2007). Characterization of reemerging chikungunya virus. PLoS Pathog 3, e89.

Stapleford, K.A., Coffey, L.L., Lay, S., Borderia, A.V., Duong, V., Isakov, O., Rozen-Gagnon, K., Arias-Goeta, C., Blanc, H., Beaucourt, S., et al. (2014). Emergence and transmission of arbovirus evolutionary intermediates with epidemic potential. Cell Host Microbe 15, 706–716.

Sun, S., Xiang, Y., Akahata, W., Holdaway, H., Pal, P., Zhang, X., Diamond, M.S., Nabel, G.J., and Rossmann, M.G. (2013). Structural analyses at pseudo atomic resolution of Chikungunya virus and antibodies show mechanisms of neutralization. Elife 2, e00435.

Tsetsarkin, K.A., Chen, R., Yun, R., Rossi, S.L., Plante, K.S., Guerbois, M., Forrester, N., Perng, G.C., Sreekumar, E., Leal, G., et al. (2014). Multi-peaked adaptive landscape for chikungunya virus evolution predicts continued fitness optimization in Aedes albopictus mosquitoes. Nat Commun 5, 4084.

Tsetsarkin, K.A., McGee, C.E., and Higgs, S. (2011). Chikungunya virus adaptation to Aedes albopictus mosquitoes does not correlate with acquisition of cholesterol dependence or decreased pH threshold for fusion reaction. Virol J 8, 376.

Tsetsarkin, K.A., Vanlandingham, D.L., McGee, C.E., and Higgs, S. (2007). A single mutation in chikungunya virus affects vector specificity and epidemic potential. PLoS Pathog 3, e201.

Tsetsarkin, K.A., and Weaver, S.C. (2011). Sequential adaptive mutations enhance efficient vector switching by Chikungunya virus and its epidemic emergence. PLoS Pathog 7, e1002412.

van Duijl-Richter, M.K., Hoornweg, T.E., Rodenhuis-Zybert, I.A., and Smit, J.M. (2015). Early Events in Chikungunya Virus Infection-From Virus Cell Binding to Membrane Fusion. Viruses 7, 3647–3674.

Vashishtha, M., Phalen, T., Marquardt, M.T., Ryu, J.S., Ng, A.C., and Kielian, M. (1998). A single point mutation controls the cholesterol dependence of Semliki Forest virus entry and exit. J Cell Biol 140, 91–99.

Vazeille, M., Moutailler, S., Coudrier, D., Rousseaux, C., Khun, H., Huerre, M., Thiria, J., Dehecq, J.S., Fontenille, D., Schuffenecker, I., et al. (2007). Two Chikungunya isolates from the outbreak of La Reunion (Indian Ocean) exhibit different patterns of infection in the mosquito, Aedes albopictus. PLoS One 2, e1168.

Vignuzzi, M., and Higgs, S. (2017). The Bridges and Blockades to Evolutionary Convergence on the Road to Predicting Chikungunya Virus Evolution. Annu Rev Virol 4, 181–200.

Voss, J.E., Vaney, M.C., Duquerroy, S., Vonrhein, C., Girard-Blanc, C., Crublet, E., Thompson, A., Bricogne, G., and Rey, F.A. (2010). Glycoprotein organization of Chikungunya virus particles revealed by X-ray crystallography. Nature 468, 709–712.

Wilder-Smith, A., Gubler, D.J., Weaver, S.C., Monath, T.P., Heymann, D.L., and Scott, T.W. (2017). Epidemic arboviral diseases: priorities for research and public health. Lancet Infect Dis 17, e101–e106.

York, A.G., Williams, K.J., Argus, J.P., Zhou, Q.D., Brar, G., Vergnes, L., Gray, E.E., Zhen, A., Wu, N.C., Yamada, D.H., et al. (2015). Limiting Cholesterol Biosynthetic Flux Spontaneously Engages Type I IFN Signaling. Cell 163, 1716–1729.

Yuan, L., Huang, X.Y., Liu, Z.Y., Zhang, F., Zhu, X.L., Yu, J.Y., Ji, X., Xu, Y.P., Li, G., Li, C., et al. (2017). A single mutation in the prM protein of Zika virus contributes to fetal microcephaly. Science 358, 933–936.

Zheng, Y., Sanchez-San Martin, C., Qin, Z.L., and Kielian, M. (2011). The domain I-domain III linker plays an important role in the fusogenic conformational change of the alphavirus membrane fusion protein. J Virol 85, 6334–6342.

Zidovetzki, R., and Levitan, I. (2007). Use of cyclodextrins to manipulate plasma membrane cholesterol content: evidence, misconceptions and control strategies. Biochim Biophys Acta 1768, 1311–1324.

