## Supplemental Figures for "Evolution-driven attenuation of alphaviruses highlights key glycoprotein determinants regulating viral infectivity and dissemination"

**Supplemental Information**

**Supplemental Figure 1. E1-V80 mutagenesis approach. (**A) Schemes showing the Sanger sequencing coverage for each experiment after 3 passages in either BHK-21 or C6/36 cells are shown. Coverages: complete structural polyprotein region (~7.6 to 11.8 Kb) and the tip of domain II (~9.9 to 10.8 Kb). (B-C) Sequence stability and acquisition of second site mutations for each mutant bearing specific E1-V80X substitution is indicated. R1 to R3 highlight individual replicates. Data is organized in stable (B) or non-stable (C) variants after BHK-21 passaging. Mutations at position 80 and second site mutations are indicated. Mutations E1-A129V/M and E1-V226A are indicated in blue and red, respectively.

**Supplemental Figure 2.** (A) Top panel. Schematic representation of the reporter viruses used in this paper. The ZsGreen and the Luciferase cassettes are depicted in dark and light gray, respectively. The circle indicated the cap, each arrow indicates the sub-genomic promoter, and the $A)_n$ indicates the poly-A tail. Lower panel. Schematic representation of the luciferase assay. (B) Replication of CHIKV E1-V80 mutants as measured by Luciferase activity. Left panel. BHK-21 cells were transfected with mRNA from the different reporter viruses, and luciferase activity was measured at the indicated time points. Right panel. Generation of infectious particles during the first round of transfections was monitored. Supernatants of the time points 4, 6 and 8 hours were used to infect BHK-21 cells. Infectious particles generation was addressed by the presence of luminescence after 24 hs post infection. GNN mutant was used as non-replicative control. Results are represented as mean and SEM, N=6 (ns, $p > 0.05$; two-way ANOVA with Bonferroni post-hoc test).

**Supplemental Figure 3. Analysis of E1-V80Q revertant variants.** (A) Upper panel. Plaque phenotype analysis of viruses isolated from E1-V80Q infected mosquitoes. A representative plaque phenotype of a E1-V80Q infected mosquitoes is depicted in M1. M10 and M28 states for two mutant variants that reverted their plaque phenotype during the course of infection in *Ae. aegypti* mosquitoes. Lower panel. Chromatograms showing the stability at position 80 and the acquisition of second site mutations. Viral loads of the infected mosquitoes are also shown. (B) Ribbon representation of E1 glycoprotein (PDBID: 3N42) highlighting position N20 is shown. Residues described in the domain I-domain III (DI-DIII) linker interaction network with the E1 core trimer are depicted in magenta (Zheng et al., 2011). Residue V80 is highlighted in red. (C) Ribbon representation of E1 glycoprotein (PDBID: 3N42) highlighting position 88 is shown. Position 88, 80 and 226 are depicted in blue, red and green, respectively.

**Supplemental Figure 4. CHIK E1-80 variants delay symptoms in *Ifnar*$^{-/-}$ mice.** (A) Body weight loss over the course of the infection. *Ifnar*$^{-/-}$ mice were inoculated via footpad injection with 1000 PFU of E1-V80, E1-V80L, E1-V80Q and carrier control. Mice were monitored twice a day for signs of infection. Data represent mean and standard deviation (N=4). (B) Disease score. Infected mice were monitored and inspected for signs of infection twice a day. We stablished a 1 to 5 disease score, where: 1-normal; 2-Inflamation in the site of inoculation; 3-Inflamation in the site of inoculation and less dynamic; 4-Hind limb inflammation, difficult to walk and hunched back; 5- dead or moribund.

**Supplemental Figure 5. CHIKV E1-V80L variants reduce cellular binding**. BHK-21 cells were incubated on ice in the presence of 20 mM ammonium chloride for 1 hour. ZsGreen E1-V80, V80A, and V80L variants were added to the cells at a MOI of 0.1, at 4 ºC and in the presence of 20 mM ammonium chloride for 5 or 60 mins. Then, cells were washed extensively with PBS, and RNA genomes were quantified by RT-qPCR.

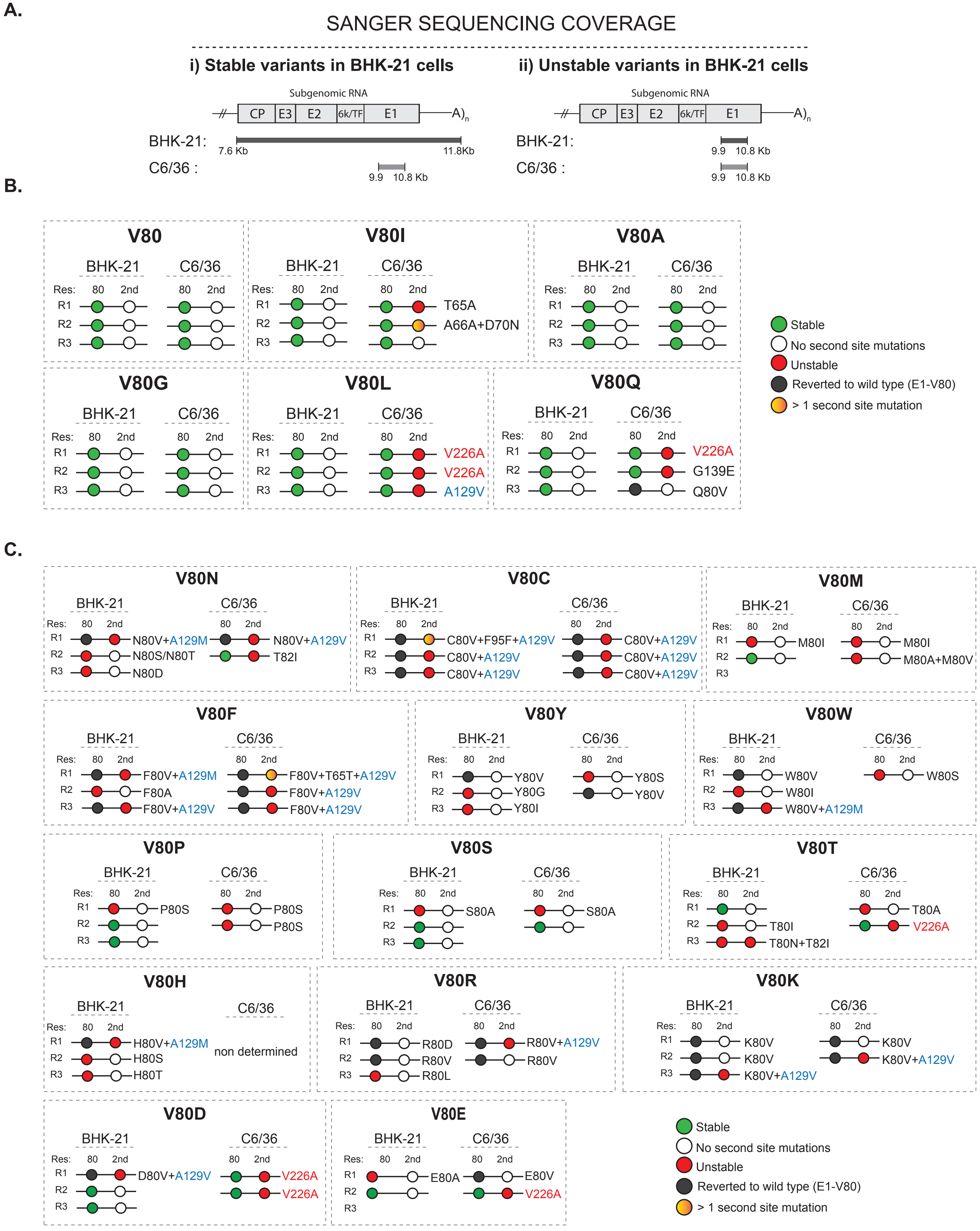


SUPPLEMENTARY FIGURE 1.

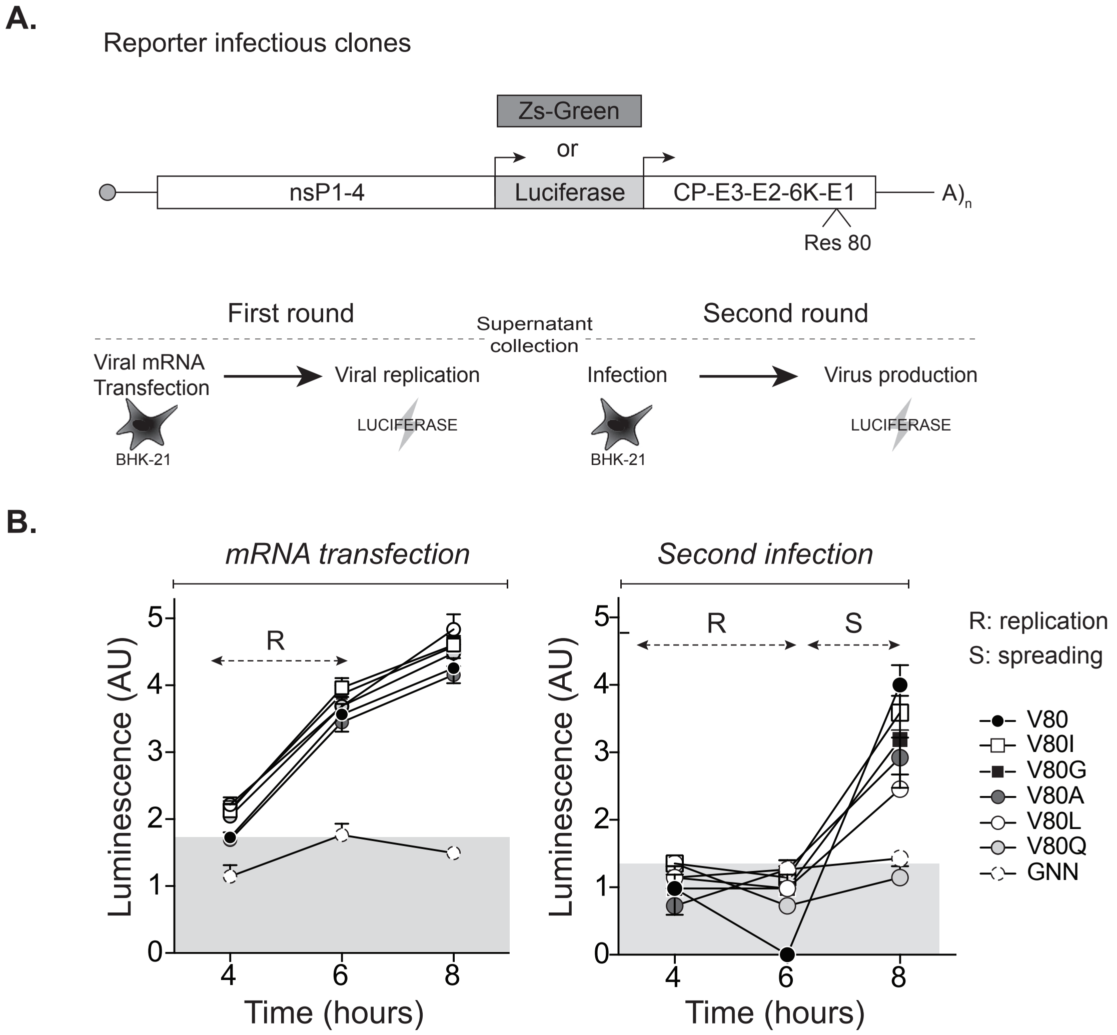


**SUPPLEMENTARY FIGURE 2.**

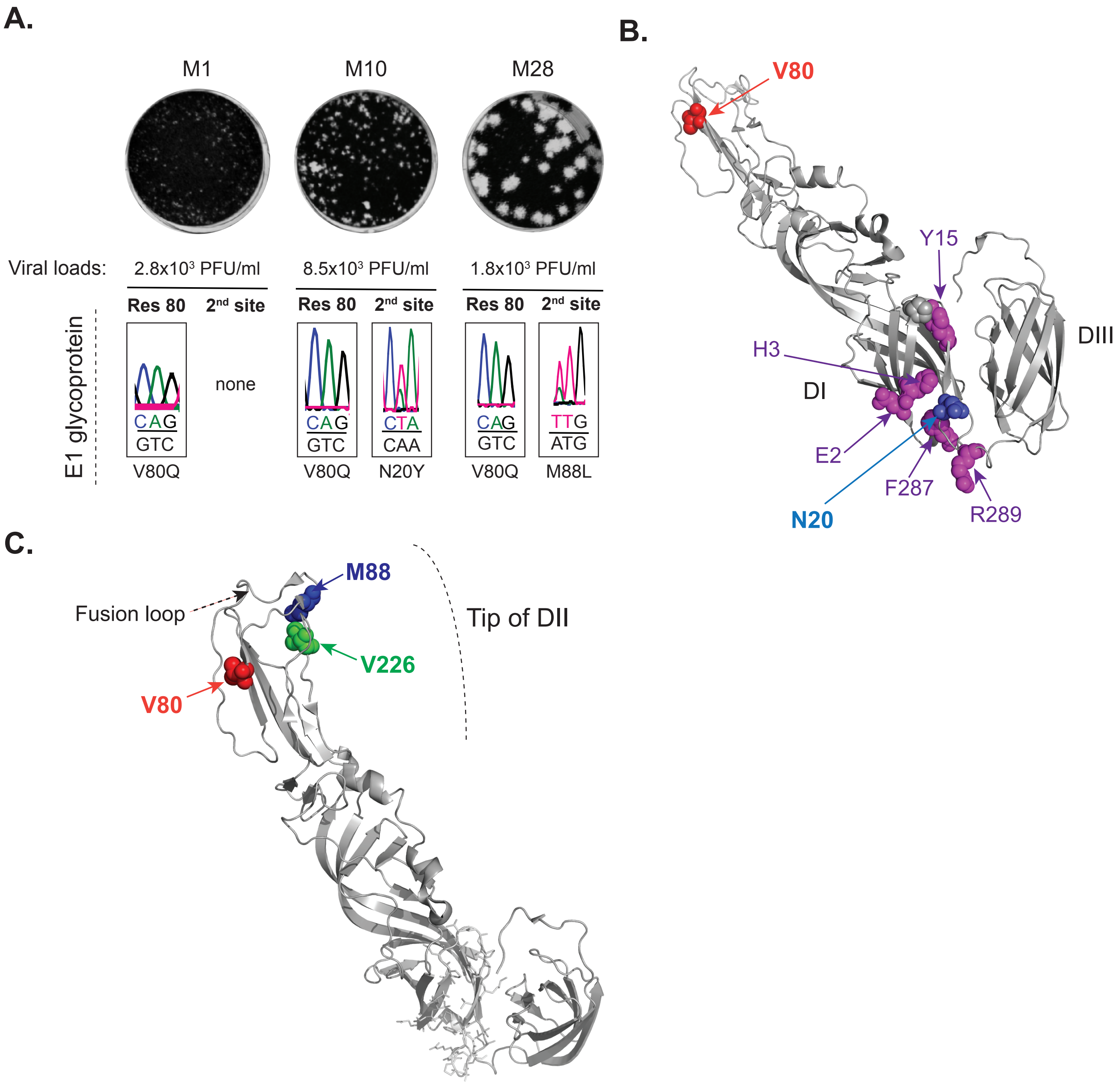


**SUPPLEMENTARY FIGURE 3.**

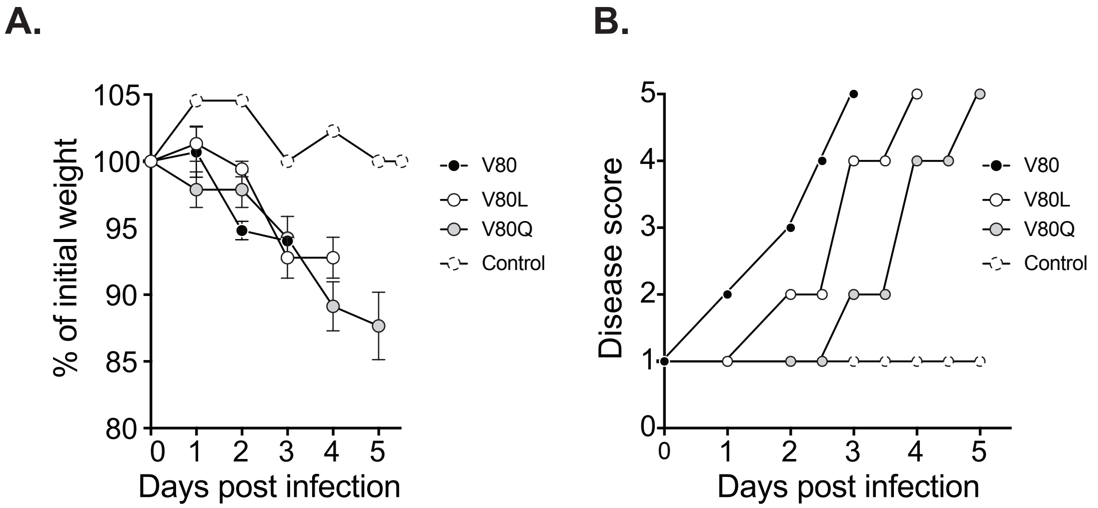


**SUPPLEMENTARY FIGURE 4.**

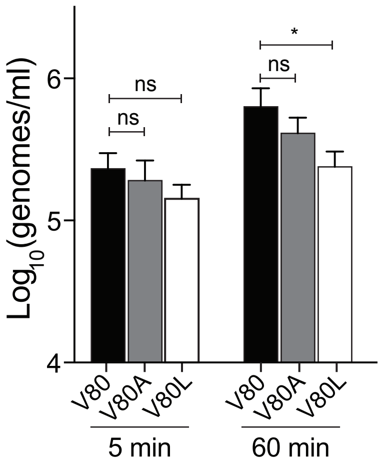


**Supplementary Figure 5**

**Supplementary Table 1**. Primers used for site directed mutagenesis and qRT-PCR

| Primer # | Name | Sequence |
|---|---|---|
| **1** | CHIKV_E1 V80R Forward | 5’-CTGACTACAGCTGTAAG**CG**CTTCACCGGCGTCTAC-3’ |
| **2** | CHIKV_E1 V80R Reverse | 5’-GTAGACGCCGGTGAAG**CG**CTTACAGCTGTAGTCAG-3’ |
| **3** | CHIKV_E1 V80H Forward | 5’-CTGACTACAGCTGTAAG**CA**CTTCACCGGCGTCTAC-3’ |
| **4** | CHIKV_E1 V80H Reverse | 5’-GTAGACGCCGGTGAAG**TG**CTTACAGCTGTAGTCAG-3’ |
| **5** | CHIKV_E1 V80K Forward | 5’-CCTGACTACAGCTGTAAG**AAG**TTCACCGGCGTCTACCC-3’ |
| **6** | CHIKV_E1 V80K Reverse | 5’-GGGTAGACGCCGGTGAA**CTT**CTTACAGCTGTAGTCAGG-3’ |
| **7** | CHIKV_E1 V80D Forward | 5’-GACTACAGCTGTAAGG**AT**TTCACCGGCGTCTACCC-3’ |
| **8** | CHIKV_E1 V80D Reverse | 5’-GGGTAGACGCCGGTGAA**AT**CCTTACAGCTGTAGTC-3’ |
| **9** | CHIKV_E1 V80E Forward | 5’-GACTACAGCTGTAAGG**AG**TTCACCGGCGTCTACCC-3’ |
| **10** | CHIKV_E1 V80E Reverse | 5’-GGGTAGACGCCGGTGAA**CT**CCTTACAGCTGTAGTC-3’ |
| **11** | CHIKV_E1 V80S Forward | 5’-CTGACTACAGCTGTAAG**TC**CTTCACCGGCGTCTAC-3’ |
| **12** | CHIKV_E1 V80S Reverse | 5’-GTAGACGCCGGTGAAG**GA**CTTACAGCTGTAGTCAG-3’ |
| **13** | CHIKV_E1 V80T Forward | 5’-CTGACTACAGCTGTAAG**AC**CTTCACCGGCGTCTAC-3’ |
| **14** | CHIKV_E1 V80T Reverse | 5’-GTAGACGCCGGTGAAG**GT**CTTACAGCTGTAGTCAG-3’ |
| **15** | CHIKV_E1 V80N Forward | 5’-CCTGACTACAGCTGTAAG**AA**CTTCACCGGCGTCTACCC-3’ |
| **16** | CHIKV_E1 V80N Reverse | 5’-GGGTAGACGCCGGTGAAG**TT**CTTACAGCTGTAGTCAGG-3’ |
| **17** | CHIKV_E1 V80Q Forward | 5’-CCTGACTACAGCTGTAAG**CAG**TTCACCGGCGTCTACCC-3’ |
| **18** | CHIKV_E1 V80Q Reverse | 5’-GGGTAGACGCCGGTGAA**CTG**CTTACAGCTGTAGTCAGG-3’ |
| **19** | CHIKV_E1 V80C Forward | 5’-CTGACTACAGCTGTAAG**TG**CTTCACCGGCGTCTAC-3’ |
| **20** | CHIKV_E1 V80C Reverse | 5’-GTAGACGCCGGTGAAG**CA**CTTACAGCTGTAGTCAG-3’ |
| **21** | CHIKV_E1 V80G Forward | 5’-GACTACAGCTGTAAGG**GA**TTCACCGGCGTCTACCC-3’ |
| **22** | CHIKV_E1 V80G Reverse | 5’-GGGTAGACGCCGGTGAA**TC**CCTTACAGCTGTAGTC-3’ |
| **23** | CHIKV_E1 V80P Forward | 5’-CTGACTACAGCTGTAAG**CC**CTTCACCGGCGTCTAC-3’ |
| **24** | CHIKV_E1 V80P Reverse | 5’-GTAGACGCCGGTGAAG**GG**CTTACAGCTGTAGTCAG-3’ |
| **25** | CHIKV_E1 V80A Forward | 5’-GACTACAGCTGTAAGG**CG**TTCACCGGCGTCTACCC-3’ |

**Supplementary Table 1**. Primers used for site directed mutagenesis and qRT-PCR

| | | |
|---|---|---|
| **26** | CHIKV_E1 V80A Reverse | 5'-GGGTAGACGCCGGTGAA**CG**CCTTACAGCTGTAGTC-3' |
| **27** | CHIKV_E1 V80L Forward | 5'-CTGACTACAGCTGTAAG**C**TCTTCACCGGCGTCTAC-3' |
| **28** | CHIKV_E1 V80L Reverse | 5'-GTAGACGCCGGTGAAGA**G**CTTACAGCTGTAGTCAG-3' |
| **29** | CHIKV_E1 V80M Forward | 5'-CCTGACTACAGCTGTAAG**ATG**TTCACCGGCGTCTACCC-3' |
| **30** | CHIKV_E1 V80M Reverse | 5'-GGGTAGACGCCGGTGAA**CAT**CTTACAGCTGTAGTCAGG-3' |
| **31** | CHIKV_E1 V80F Forward | 5'-CTGACTACAGCTGTAAG**T**TCTTCACCGGCGTCTAC-3' |
| **32** | CHIKV_E1 V80F Reverse | 5'-GTAGACGCCGGTGAAGA**A**CTTACAGCTGTAGTCAG-3' |
| **33** | CHIKV_E1 V80Y Forward | 5'-CCTGACTACAGCTGTAAG**TA**CTTCACCGGCGTCTACC-3' |
| **34** | CHIKV_E1 V80Y Reverse | 5'-GGGTAGACGCCGGTGAAG**TA**CTTACAGCTGTAGTCAGG-3' |
| **35** | CHIKV_E1 V80W Forward | 5'-CCTGACTACAGCTGTAAG**TGG**TTCACCGGCGTCTACCC-3' |
| **36** | CHIKV_E1 V80W Reverse | 5'-GGGTAGACGCCGGTGAA**CCA**CTTACAGCTGTAGTCAGG-3' |
| **37** | CHIKV_E1 V80I Forward | 5'-CCTGACTACAGCTGTAAG**A**TCTTCACCGGCGTCTACCC-3' |
| **38** | CHIKV_E1 V80I Reverse | 5'-GGGTAGACGCCGGTGAAGA**T**CTTACAGCTGTAGTCAGG-3' |
| **39** | CHIKV_E1 V226A Forward | 5'-CTGCAGAGACCGGCTG**C**GGGTACGGTACACGTG-3' |
| **40** | CHIKV_E1 V226A Reverse | 5'-CACGTGTACCGTACCC**G**CAGCCGGTCTCTGCAG-3' |
| **41** | SINV_E1(V80L) Forward | 5'-CAGACTATACCTGCAAG**C**TCTTCGGAGGGGTCTACC-3' |
| **42** | SINV_E1(V80L) Reverse | 5'-GGTAGACCCCTCCGAAGA**G**CTTGCAGGTATAGTCTG-3' |
| **43** | ONNV_E1(V80L) Forward | 5'-CCAGATTATAACTGCAAA**CTC**TTCACAGGCGTCTACCC-3' |
| **44** | ONNV_E1(V80L) Reverse | 5'-GGGTAGACGCCTGTGAA**GAG**TTTGCAGTTATAATCTGG-3' |
| **45** | CHIKV. qRT-PCR Forward | 5' TCACTCCCTGCTGGACTTGATAGA 3' |
| **46** | CHIKV. qRT-PCR Reverse | 5' TTGACGAACAGAGTTAGGAACATACC 3' |
| **47** | CHIKV. qRT-PCR Probe | 5' AGGTACGCGCTTCAAGTTCGGCG 3' |
